# Dissecting genetic variance structure and evaluating genomic prediction models for single-cross hybrids derived from Stiff Stalk and Non-Stiff Stalk maize heterotic groups

**DOI:** 10.64898/2026.03.11.710575

**Authors:** Jenifer Camila Godoy dos Santos, Jode Edwards, Elizabeth Lee, Mark A. Mikel, Samuel B. Fernandes, Candice N. Hirsch, Sydney P. Berry, Alexander E. Lipka, Martin O. Bohn

## Abstract

The early 20th-century discovery of heterosis and the establishment of heterotic groups transformed maize (*Zea mays* L.) into a keystone of global agriculture. However, maize breeding faces two significant challenges: the gradual decline of general combining ability (GCA) variance within heterotic groups and the impracticality of testing all possible single crosses in the early stages of a breeding program. Here, we developed genomic best linear unbiased prediction (GBLUP)-based multi-kernel models, using additive and two alternative non-additive genomic relationship matrices, to estimate the variance components associated with the GCA of Stiff Stalk (SS) and Non-Stiff Stalk (NSS) heterotic groups and the specific combining ability (SCA) arising from their crosses. We further applied these models to predict the performance of untested single-cross combinations under varying levels of parental information. We showed that the SS and NSS groups retained significant GCA variance across traits in both early- and late-maturity groups. The SS group, in contrast, exhibited no detectable GCA variance in grain yield for the intermediate-flowering subset of hybrids, highlighting a limitation for future genetic improvement. Furthermore, our results showed that GBLUP-based multi-kernel models effectively identified superior hybrids when parental information was available. In the absence of this information, however, these models underperformed compared to covariance-based approaches. Both types of non-additive matrices produced similar results, affirming the robustness of the inferred genetic architecture. Overall, this study sheds light on the future use of US maize commercial germplasm and demonstrates how GBLUP-based multi-kernel models can improve the efficiency of hybrid breeding programs.

## Introduction

Modern maize breeding has its roots in the early 20th century, when the discovery of heterosis revolutionized hybrid development (Shull 1908; East 1908). Since then, the establishment of heterotic groups has been central to maximizing heterosis. A heterotic group is a collection of inbred lines that are relatively similar to each other but deliver superior hybrid performance when crossed with inbred lines from a contrasting group (Melchinger and Gumber 1998). In US commercial germplasm, one of the most prominent heterotic patterns is the division between the Stiff Stalk (SS) and Non-Stiff Stalk (NSS) groups. The SS group traces its ancestry to the Iowa Stiff Stalk Synthetic population and includes inbred lines such as B37 and B73 (Troyer 1999). The NSS group, further subdivided into Iodent and non-Iodent, consists of genetically diverse sources and encompasses important inbred lines such as Mo17 and PH207 (Mikel and Dudley 2006).

At its core, maize breeding is a long-term reciprocal recurrent selection program, where two heterotic groups and their intercrosses are improved simultaneously. In practice, this process generally involves crossing inbred lines from one group with testers from the opposite group to estimate the general combining ability (GCA) of the inbred lines and the specific combining ability (SCA) of the resulting hybrids (Sprague and Tatum 1942). The GCA can reflect additive effects and indicates how consistently a parental line contributes to superior hybrid performance across various crosses. By estimating GCA, breeders can identify the most promising inbred lines to be selected and recombined, thereby promoting the accumulation of favorable alleles and progressively enhancing the genetic potential of each heterotic group. At the same time, the preferential selection and eventual fixation of different alleles across groups increase their genetic distance, reinforcing the heterotic pattern. In contrast, SCA captures the hybrid’s heterotic response due to non-additive effects that frequently involve dominance and/or epistasis (Garcia *et al*. 2025). A significant SCA indicates that the hybrid confers an advantageous performance that cannot be explained solely by the GCA of its parents. Thus, hybrid performance is determined by the combined contribution of parental GCA and their SCA, and the joint evaluation of both is crucial for identifying superior hybrids and advancing them toward commercial release (Comstock et al. 1949; Bernardo *et al*. 2002).

A significant challenge in maize breeding is the gradual loss of additive genetic variance within heterotic groups, which occurs due to the high intensity of selection implemented over multiple breeding cycles (Hallauer 1992). This issue is particularly pronounced in commercial heterotic groups with a narrow genetic base, such as the SS group, which was established from a limited number of founder lines (Troyer 1999). Although these lines still retain a considerable amount of maize’s genetic diversity, the historical dependence on a small set of ancestors has rendered the SS group more susceptible to the erosion of additive genetic variance over time. If not managed effectively, this vulnerability can lead to a decline in GCA within this group, ultimately restricting the genetic progress possible in inter-group hybrids in the long term (Goodman 2005).

A second challenge faced by maize breeders is the impracticability of evaluating all possible crosses between inbred lines, particularly in the early stages of a program. To address this limitation, several strategies have been developed to evaluate the performance of hybrids that have not yet been field-tested. An early approach was proposed by Bernardo (1994), who used RFLP markers to estimate genetic relationships among parental lines and incorporated this information to predict the untested crosses. With the advent of high-density genotyping, this concept was expanded through genomic selection (GS), which exploits genome-wide marker data to model the genetic merit of individuals before any phenotypic evaluation (Meuwissen et al. 2001). In the context of maize breeding, GS enables breeders to assess the entire set of possible single-cross hybrids solely from parental genomic information. This capability, in turn, allows the identification of promising combinations and the filtering out of those with low potential without the need for extensive phenotypic testing (Smith et al. 2004; Technow et al. 2014; Kadam et al. 2016).

Among GS models, genomic best linear unbiased prediction (GBLUP) is one of the most widely applied in maize breeding (Cantelmo et al. 2017; Dalsente Krause et al. 2020). In its standard form, the GBLUP model relies on an additive relationship matrix derived from molecular markers to make predictions (VanRaden 2008). By accounting for realized genetic resemblance between individuals, GBLUP eliminates the bias that can occur in pedigree-based models, which assume that all close relatives share the same level of genetic relationship (Gamal El-Dien et al. 2016). Furthermore, GBLUP can be extended to multi-kernel models, in which non-additive relationship matrices are incorporated alongside the additive matrix to also account for variation arising from dominance and epistasis effects (Su et al. 2012; Vitezica et al. 2013). The addition of these matrices can boost the model’s predictive ability and provide breeders with a deeper understanding of the genetic architecture of the traits being studied (Dias et al. 2018; Peixoto et al. 2024). These matrices also ensure an orthogonal separation of additive, dominance, and epistatic effects, which facilitates a clearer distinction between genetic variance components (Muñoz et al. 2014; Nazarian and Gezan 2016).

A set of 162 single-cross hybrids derived from SS and NSS inbred lines using a North Carolina design II mating scheme (Comstock and Robinson 1948) is considered for this study. We developed GBLUP-based multi-kernel models, using additive and non-additive genomic relationship matrices, to partition the total phenotypic variability into variance components associated with the GCA of SS and NSS lines, as well as the non-additive variance captured by SCA. For SCA, two alternative non-additive matrices were tested, and different variance–covariance structures were specified to explicitly model genotype-by-environment (GEI) effects. Using these models, we also predicted the performance of untested single-cross combinations across various training set configurations, which ranged from full parental representation to none. Our work addressed two fundamental aspects that could have major implications for maize breeding. First, we examined whether the SS and NSS parental lines included in this study, which represent the long-established SS and NSS heterotic groups, still harbor sufficient GCA variance to sustain future gains. Second, we evaluated to what extent GBLUP-based multi-kernel models, which simultaneously integrate additive, non-additive, and GEI effects, enhance predictive ability and yield interpretable insights into the architecture of hybrid performance under varying levels of parental information.

## Materials and methods

### Genetic material

A diverse set of inbred lines from the two major maize heterotic groups of the US Corn Belt, SS and NSS, was assembled to serve as the parental pool for this study (Troyer 1999). The set comprised of 13 SS and 28 NSS lines, selected for their relevance to the commercial maize breeding industry.

To maximize genetic diversity and ensure proper synchronization of flowering times, the inbred lines were categorized into early, intermediate, and late maturity groups (Table 1). A North Carolina design II mating scheme (Comstock and Robinson 1948) was followed to generate single-cross hybrids between SS lines (seed parent) and NSS lines (pollen parent). While 72 early, 70 intermediate, and 75 late hybrids were theoretically possible, a total of 40, 67, and 74 hybrids were produced, respectively. The shortfall was primarily due to practical constraints, including flowering asynchrony and limited seed availability. Since some inbred lines were assigned to more than one maturity group, the total number of unique single-cross hybrids was 162, which is fewer than the total number of hybrids across the three groups (Supplementary Tables S1-S4).

**Table 1.** Stiff Stalk (SS) and Non-Stiff Stalk (NSS) inbred lines assigned to the early, intermediate, and late maturity groups according to their relative maturity classification.

| Early |  | Intermediate |  | Late |  |
| --- | --- | --- | --- | --- | --- |
| SS | NSS | SS | NSS | SS | NSS |
| LH145 | PHR25 | PHN66 | PHG83 | PHHB9 | PHN47 |
| CG120 | PHN82 | PHB47 | Q381 | PHV63 | Q381 |
| PHRE1 | PHG29 | PHW52 | PHN82 | PHP38 | PHN82 |
| PHB47 | LH82 | LH198 | PHG29 | LH195 | PHW30 |
| PHHV4 | PHZ51 | LH195 | PHW30 | PHW52 | PHR63 |
| S8324 | LH185 |  | LH82 |  | PHM49 |
|  | PHK56 |  | PHZ51 |  | PHW53 |
|  | PHG47 |  | PHR55 |  | PHZ51 |
|  | LH162 |  | LH185 |  | PHR55 |
|  | CG123 |  | PHK76 |  | PHR03 |
|  | PHW03 |  | LH38 |  | LH185 |
|  | PHK05 |  | PHK56 |  | PHP60 |
|  |  |  | PHG47 |  | LH210 |
|  |  |  | LH51 |  | PHJ65 |
|  |  |  |  |  | PHM57 |

### Phenotypic data

As part of the Genomes to Fields (G2F) initiative (McFarland *et al*. 2020), the 162 single-cross hybrids were evaluated in multilocation field trials. In 2016, they were tested across 27 US locations and 2 Canadian locations, and in 2017, across 29 US locations and 2 Canadian locations. In both years, these hybrids were assessed alongside additional hybrids not included in this study but present in the same trials. These locations were grouped into three maturity sets, namely early, intermediate, and late, with an unbalanced distribution of individuals both within and between years. For analytical purposes, each unique combination of year and location was considered a distinct environment, resulting in a total of 60 environments. In each environment, trials were conducted using a randomized complete block design with two blocks, except in one instance where four blocks were used.

Although several agronomic traits were initially evaluated in the G2F trials, we focused on the following traits for our down-stream analyses: grain yield (GY), reported in tons per hectare (t ha^−1^), plant height (PH) and ear height (EH), measured in centimeters; silking (SI), defined as the number of days from planting until silk emergence; and anthesis (AN), defined as the number of days from planting until pollen shed. These traits were assessed according to the standard operating procedures established by the G2F initiative (Genomes to Fields Initiative 2025).

To ensure high-quality and consistent phenotypic information, we applied an initial round of filtering to the entire dataset. This filtering included not only the hybrids of interest but also all hybrids evaluated in the G2F trials. Thus, the final data set included 806 maize hybrids and 44 environments. Importantly, none of the early, intermediate, and late single crosses needed to be removed during this process and are included among the 806 hybrids.

### Phenotypic analyses

#### Single-environment trial analyses

The following model was fitted to the entire filtered phenotypic data set to perform a single-environment trial analysis for each trait:

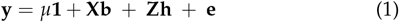

where **y** (*n* × 1) is the vector of phenotypic observations with *n* plots resulting from the combination of *g* hybrids and *r* blocks; *µ* is the intercept; **b** (*r* × 1) is the vector of fixed effects of blocks; **h** (*g* × 1) is the vector of random effects of hybrids, with 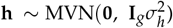 and **e** (*n* × 1) is the vector of random effects of residuals, with 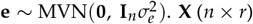 and **Z** (*n* × *g*) represent incidence matrices for their respective effects, **1** (*n* × 1) denotes a vector of ones, **I**_*g*_ and **I**_*n*_ indicate identity matrices of appropriate dimensions, and 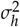 and 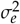 denote the variance components of hybrids and residuals, respectively. Based on this model, we calculated the broad-sense heritability (*H*^2^) (Cullis et al. 2006) and the coefficient of variation (*CV*), as follows:

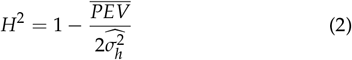

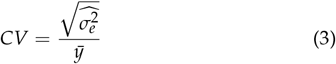

where 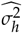 and 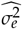 are the estimated variance components of hybrids and residuals, respectively; 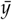 is the mean trait value within each environment; and 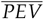 denotes the mean prediction error variance of the difference between two empirical best linear unbiased predictions (E-BLUPs) of the hybrid effects. The 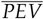 represents the mean uncertainty associated with pairwise comparisons among hybrids and is therefore central to the definition of Cullis heritability (Cullis et al. 2006).

We also fitted a model similar to (1), this time treating hybrids as fixed effects, to obtain their empirical best linear unbiased estimations (E-BLUEs). Subsequently, we excluded environments with *CV* greater than 25% and/or those with near-zero *H*^2^ from this set of E-BLUEs (Supplementary Table S5). In addition, we retained only the E-BLUEs corresponding to hybrids from the early, intermediate, and late single-cross groups. The resulting filtered data set of E-BLUEs for 162 single-cross hybrids at 37 environments was used as the response variable in all subsequent analyses. A summary of the phenotypic distributions and trait correlations, along with the co-occurrence matrix of hybrids across environments, is presented in Figures S1 and S2. Table S6 further provides the classification of these environments into maturity groups and their geographical distribution.

#### Multi-environment trial analyses

For each trait, we also performed a multi-environment trial analysis by fitting the following linear mixed model:

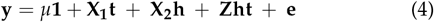

where **y** (*n* × 1) is the vector of E-BLUEs for *s* = 37 environments and *g* = 162 single-cross hybrids; 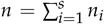, with *n*_*i*_ being the number of single-cross hybrids in environment *i*; *µ* is the intercept; **1** (*n* × 1) is a vector of ones; **t** (*s* × 1) is the vector of fixed effects of environments; and **h** (*g* × 1) is the vector of fixed effects of hybrids. The term **ht** (*gs* × 1) represents the random interaction between hybrids and environments, modeled as **ht** ∼ MVN (**0, G**_*gs*_), where 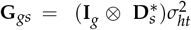. Here, **I**_*g*_ is an identity matrix of proper dimension, ⨂ denotes the Kronecker product, and 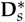 is a diagonal matrix whose entries correspond to the environment-specific variance component, 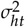, associated with the interaction effects. **X**_**1**_ (*n* × *s*), **X**_**2**_ (*n* × *g*), and **Z** (*n* × *gs*) are incidence matrices for their respective effects. In this framework, the variance-covariance matrix for the residual term **e** (*n* × 1) was predefined as a diagonal matrix, with elements specified as:

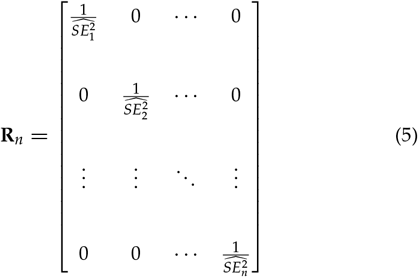

where 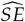 terms denote the standard errors derived from the single-trial analysis (Frensham *et al*. 1997). Using this multi-environmental model, we obtained E-BLUEs for hybrids, which were used in genomic prediction analyses.

### Genotypic data

The SS and NSS lines belong to a broader panel of inbred lines from the G × E Project of the G2F initiative (Lima *et al*. 2023), which employed the Practical Haplotype Graph platform to generate variant calls (Bradbury *et al*. 2022). From this complete data set, we used VCFtools v0.1.15 (Danecek *et al*. 2011) to select only the individuals of interest, resulting in a subset of 41 inbred lines (13 SS and 28 NSS) and 437,214 SNPs.

For quality control, we used VCFtools v0.1.15 to filter out SNPs that were not bi-allelic, had a minor allele frequency (MAF) below 5%, or exhibited more than 20% missing genotypes. We also removed redundant markers in high linkage disequilibrium (LD) using the LD pruning algorithm available in PLINK v1.9 (Purcell *et al*. 2007). This algorithm employed a sliding window approach to evaluate squared Pearson’s correlation coefficient (*r*^2^) among SNPs within windows of 100 markers. When a pair of SNPs within a window displayed an *r*^2^ value greater than 0.9, one of them was removed. The window was then advanced by 20 markers, and the procedure repeated. This approach preserved only one representative SNP from each highly correlated region, resulting in a final data set of 112,199 SNPs.

After applying quality control filters, we inferred the genotypes of all possible single-cross hybrids derived from the combinations between the 13 SS and 28 NSS parental lines using TASSEL v5 (Bradbury et al. 2007). Finally, we converted the genotypic data of both the parental lines and the inferred hybrids into a numeric format using the simplePHENOTYPES R package v1.3.0 (Fernandes and Lipka 2020).

### Genomic relationship matrices

Following the methodology proposed by VanRaden (2008), we computed the additive genomic relationship matrix (**A**) using the filtered and converted genotypic data of the 41 parental lines, based on:

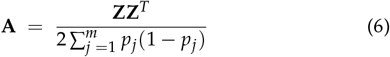

with:

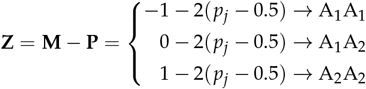

where **M** (*n*_*g*_ × *m*) is a marker matrix centered by the MAF; **P** (*n*_*g*_ × *m*) is a matrix of allele frequencies, with each column representing a locus and being expressed as *P*_*j*_ = 2(*p*_*j*_ − 0.5); *p*_*j*_ is the MAF of the *j*^*th*^ locus; *n*_*g*_ is the number of genotyped individuals; and *m* is the total number of markers. The matrix **A**, which represents the covariance structure for GCA effects, was then divided into **A**_**SS**_, describing the relationships among the 13 SS lines, and **A**_**NSS**_, describing the relationships among the 28 NSS lines.

To model the covariance structure of SCA effects in the single-cross hybrids, we explored two alternative strategies. The first approach involved the construction of a non-additive genomic relationship matrix (**D**) from the inferred genotypes, according to the method of Vitezica *et al*. (2013):

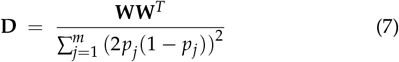

with:

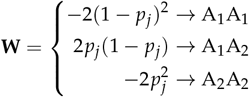

where all terms were previously defined. In both **A** and **D** matrices, we assumed that A_2_ corresponded to the least frequent allele, resulting in genotype values of 0, 1, and 2 for the genotypes A_1_A_1_ (major homozygote), A_1_A_2_ (heterozygote), and A_2_A_2_ (minor homozygote), respectively.

The second approach relied on a formulation described by Stuber and Cockerham (1966), which uses additive genetic relationships between parental lines to construct a covariance matrix, referred to here as the matrix **S**. Specifically, for a given pair of single crosses, H_*kl*_ = SS_*k*_ × NSS_*l*_ and H_*k*′ *l*′_ = SS_*k*′_ × NSS_*l*′_, the corresponding element of **S** was defined as:

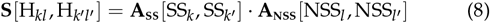

where SS_*k*_ and SS_*k*′_ denote the *k*^*th*^ and *k*′^*th*^ SS lines, with *k, k*′ ∈ 1, …, *a* and *a* = 13. Similarly, NSS_*l*_ and NSS_*l*′_ denote the *l*^*th*^ and *l*^′*th*^ NSS lines, with *l, l*^′^ ∈ 1, …, *b* and *b* = 28. The terms **A**_**SS**_[SS_*k*_, SS_*k*′_ ] and **A**_**NSS**_[NSS_*l*_, NSS_*l*′_ ] correspond to the respective entries of the **A**_**SS**_ and **A**_**NSS**_ matrices, with the dot (·) indicating multiplication between them. It is worth noting that both **D** and **S** were built considering all possible single-cross hybrids derived from the combinations between the 13 SS and 28 NSS parental lines.

The **A** and **D** matrices were calculated using the AGHmatrix R package v2.1.4 (Amadeu *et al*. 2016, 2023), whereas the **S** matrix was derived from a custom R function developed from the theoretical foundation described above. Because some of these matrices were not positive definite, their inverses were obtained using iterative bending methods, as implemented in the ASRgenomics package v1.1.4 (Gezan *et al*. 2022).

### GBLUP-based multi-kernel models

#### Model specifications

A set of GBLUP-based multi-kernel models was fitted for each trait, taking on the following form:

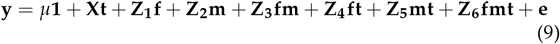

where **y** (*n* × 1) is the vector of E-BLUEs for *s* = 37 environments and *g* = 162 single-cross hybrids; 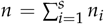, with *n*_*i*_ being the number of single-cross hybrids in environment *i*; *µ* is the intercept; **1** (*n* × 1) is a vector of ones; and **t** (*s* × 1) is the vector of fixed effects of environments with incidence matrix **X** (*n* × *s*). The vectors **f** (*a* × 1), **m** (*b* × 1), and **fm** (*ab* × 1) represent, respectively, the GCA effects of seed parents (SS lines), the GCA effects of pollen parents (NSS lines), and the SCA effects arising from their interaction, with corresponding incidence matrices **Z**_**1**_ (*n* × *a*), **Z**_**2**_ (*n* × *b*), and **Z**_**3**_ (*n* × *ab*). The interactions between seed and pollen parents with environments are represented by the vectors **ft** (*as* × 1) and **mt** (*bs* × 1), along with their respective incidence matrices **Z**_4_ (*n* × *as*) and **Z**_5_ (*n* × *bs*). The vector **fmt** (*abs* × 1), with incidence matrix **Z**_**6**_ (*n* × *abs*), accounts for the three-way interaction among seed parents, pollen parents, and environments. All vectors **f, m, fm, ft, mt, fmt**, and the residual **e** were treated as random and assumed to be independent and normally distributed with zero means and variance–covariance matrices equal to:

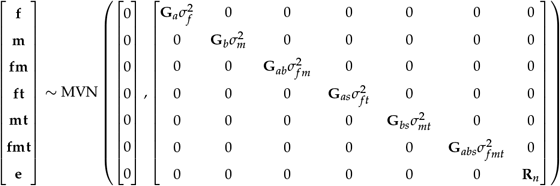

where each *σ*^2^ term denotes the variance components of the random effects and **R**_*n*_ denotes the residual variance–covariance matrix, as specified in (5). The variance–covariance matrices **G** were modeled using alternative structures, resulting in a collection of model configurations. The differences among these models pertain to the specification of **G**_*ab*_, modeled using either the inverse of the **D** or the **S** matrix, and of **G**_*as*_, **G**_*bs*_, and **G**_*abs*_, which were specified using either an identity matrix (**I**) or a diagonal matrix 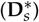 (see Table 2).

**Table 2.** Summary of the six GBLUP-based multi-kernel models considered for each trait, including variance–covariance structures for parental effects, their interactions, and environment-specific components.

| Model | Variance–Covariance Structure |  |  |  |  |  |
| --- | --- | --- | --- | --- | --- | --- |
| | $\mathbf{G}_a$ | $\mathbf{G}_b$ | $\mathbf{G}_{ab}$ | $\mathbf{G}_{as}$ | $\mathbf{G}_{bs}$ | $\mathbf{G}_{abs}$ |
| M1-D | $\mathbf{A}_{SS}$ | $\mathbf{A}_{NSS}$ | $\mathbf{D}$ | $\mathbf{I}_a \otimes \mathbf{I}_s$ | $\mathbf{I}_b \otimes \mathbf{I}_s$ | $\mathbf{I}_a \otimes \mathbf{I}_b \otimes \mathbf{I}_s$ |
| M2-D | $\mathbf{A}_{SS}$ | $\mathbf{A}_{NSS}$ | $\mathbf{D}$ | $\mathbf{I}_a \otimes \mathbf{I}_s$ | $\mathbf{I}_b \otimes \mathbf{I}_s$ | $\mathbf{I}_a \otimes \mathbf{I}_b \otimes \mathbf{D}_s^*$ |
| M3-D | $\mathbf{A}_{SS}$ | $\mathbf{A}_{NSS}$ | $\mathbf{D}$ | $\mathbf{I}_a \otimes \mathbf{D}_s^*$ | $\mathbf{I}_b \otimes \mathbf{D}_s^*$ | $\mathbf{I}_a \otimes \mathbf{I}_b \otimes \mathbf{D}_s^*$ |
| M1-S | $\mathbf{A}_{SS}$ | $\mathbf{A}_{NSS}$ | $\mathbf{S}$ | $\mathbf{I}_a \otimes \mathbf{I}_s$ | $\mathbf{I}_b \otimes \mathbf{I}_s$ | $\mathbf{I}_a \otimes \mathbf{I}_b \otimes \mathbf{I}_s$ |
| M2-S | $\mathbf{A}_{SS}$ | $\mathbf{A}_{NSS}$ | $\mathbf{S}$ | $\mathbf{I}_a \otimes \mathbf{I}_s$ | $\mathbf{I}_b \otimes \mathbf{I}_s$ | $\mathbf{I}_a \otimes \mathbf{I}_b \otimes \mathbf{D}_s^*$ |
| M3-S | $\mathbf{A}_{SS}$ | $\mathbf{A}_{NSS}$ | $\mathbf{S}$ | $\mathbf{I}_a \otimes \mathbf{D}_s^*$ | $\mathbf{I}_b \otimes \mathbf{D}_s^*$ | $\mathbf{I}_a \otimes \mathbf{I}_b \otimes \mathbf{D}_s^*$ |

$\mathbf{A}_{SS}$ : additive genomic relationship matrix for the SS inbred lines; $\mathbf{A}_{NSS}$ : additive genomic relationship matrix for the NSS inbred lines; $\mathbf{D}$ : non-additive genomic relationship matrix for all possible single-cross hybrids derived from the combinations of the parental lines; $\mathbf{S}$ : covariance matrix obtained from additive genetic relationships between parental lines; $\mathbf{I}_a$ , $\mathbf{I}_b$ , $\mathbf{I}_s$ : identity matrices assuming homogeneous variances across environments; $\mathbf{D}_s^*$ : diagonal matrix assuming a specific variance for each environment; and $\otimes$ : Kronecker product. The subscripts $a$ , $b$ , and $s$ indicate matrix dimensions, consistent with the definitions provided earlier in the text.

To compare the models, we assessed their goodness of fit using the Akaike Information Criterion (AIC) (Akaike 1974), defined as:

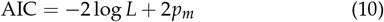

where *L* is the maximum point of the residual likelihood function and *p*_*m*_ is the number of estimated parameters. Considering the alternative variance–covariance structures, we retained the model with the lowest AIC among those specified with the **D** matrix and, independently, the one with the lowest AIC among those specified with the **S** matrix. We also used diagnostic plots to identify potential outliers and evaluate the normality of residuals in the chosen models.

#### Estimation of genetic variance components

Using the best-fitting models defined by the **D** and **S** matrices (Table 2), we estimated the variance components associated with the GCA of parental lines and the non-additive variance captured by the SCA of their interaction. To evaluate their significance, we conducted likelihood ratio tests (LRT), under the assumption that the statistic follows a mixed chi-square distribution (Self and Liang 1987). This process involved comparing the full model with the reduced models, where each reduced model was obtained by excluding the variance component of interest. Furthermore, we applied the best-fitting models separately for each maturity group to obtain group-specific variance component estimates. In these analyses, environments with limited representation of parental lines and hybrids were excluded to minimize confounding effects, resulting in 18, 36, and 28 environments retained in the early, intermediate, and late groups, respectively.

#### Genomic prediction of single-cross hybrids

We tested four methods to predict the performance of untested single-cross hybrids 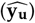 (Table 3). In the first two methods, predictions were generated using only the GCA effects of the parental lines and combining the GCA effects with the SCA (described in Technow *et al*. (2014) and Kadam *et al*. (2016)):

**Table 3.** Genomic prediction methods used to infer the performance of untested single-cross hybrids.

| Method | Variant | Prediction equation | Source of components |
| --- | --- | --- | --- |
| 1 | a | $\hat{\mathbf{y}}_{\mathbf{u}} = \hat{\mu} + \hat{\mathbf{f}} + \hat{\mathbf{m}}$ | multi-kernel model with **D** |
| | b | $\hat{\mathbf{y}}_{\mathbf{u}} = \hat{\mu} + \hat{\mathbf{f}} + \hat{\mathbf{m}}$ | multi-kernel model with **S** |
| 2 | a | $\hat{\mathbf{y}}_{\mathbf{u}} = \hat{\mu} + \hat{\mathbf{f}} + \hat{\mathbf{m}} + \hat{\mathbf{f}}\mathbf{m}$ | multi-kernel model with **D** |
| | b | $\hat{\mathbf{y}}_{\mathbf{u}} = \hat{\mu} + \hat{\mathbf{f}} + \hat{\mathbf{m}} + \hat{\mathbf{f}}\mathbf{m}$ | multi-kernel model with **S** |
| 3 | a | $\hat{\mathbf{y}}_{\mathbf{u}} = \mathbf{C}_{\mathbf{ut}}\mathbf{C}_{\mathbf{tt}}^{-1}\hat{\mathbf{y}}$ | **A<sub>SS</sub>** and **A<sub>NSS</sub>**; multi-kernel model with **D** |
| | b | $\hat{\mathbf{y}}_{\mathbf{u}} = \mathbf{C}_{\mathbf{ut}}\mathbf{C}_{\mathbf{tt}}^{-1}\hat{\mathbf{y}}$ | **A<sub>SS</sub>** and **A<sub>NSS</sub>**; multi-kernel model with **S** |
| 4 | a | $\hat{\mathbf{y}}_{\mathbf{u}} = \mathbf{C}_{\mathbf{ut}}\mathbf{C}_{\mathbf{tt}}^{-1}\hat{\mathbf{y}}$ | **A<sub>SS</sub>**, **A<sub>NSS</sub>**, and **D**; multi-kernel model with **D** |
| | b | $\hat{\mathbf{y}}_{\mathbf{u}} = \mathbf{C}_{\mathbf{ut}}\mathbf{C}_{\mathbf{tt}}^{-1}\hat{\mathbf{y}}$ | **A<sub>SS</sub>**, **A<sub>NSS</sub>**, and **S**; multi-kernel model with **S** |

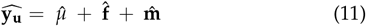

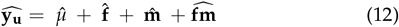

where 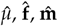, and 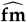 were estimated from the best-fitting GBLUP-based multi-kernel models (Table 2). Even in method 1 (presented in Equation 11), two variants were needed because GCA estimates differed depending on whether SCA was specified with **D** or **S** in model 9.

For comparison purposes, the last two methods were based on the covariance between tested and untested single-cross hybrids, as outlined in Bernardo (1996):

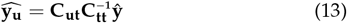

where **C**_**ut**_ and **C**_**tt**_ represent the genetic covariance matrices between untested and tested, and among tested single crosses, respectively, and **ŷ** denotes the vector of E-BLUEs for the tested hybrids, obtained from model 4. In method 3, the elements of **C**_**ut**_ and the off-diagonal elements of **C**_**tt**_ for a given pair of single crosses were computed as:

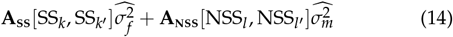

where all terms retain their previously defined meaning. The diagonal elements of **C**_**tt**_ were calculated as:

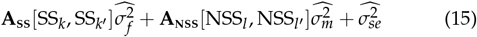

where 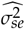 was obtained as the square of the standard errors associated with the E-BLUEs estimated from Equation 4. As in method 1, in method 3 the variance components were taken from the best-fitting **D** and **S** models, giving rise to variants 3a and 3b. This comes from the fact that all variance components are jointly estimated within the context of a mixed model, and their values are influenced by the specification of the full covariance structure (Henderson 1975). In method 4, these covariance matrices were extended to incorporate hybrid-level information derived from non-additive genomic relationships. Accordingly, the elements of **C**_**ut**_ and the off-diagonal elements of **C**_**tt**_ were computed as:

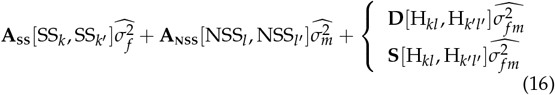

and the diagonal elements of **C**_**tt**_ were calculated as:

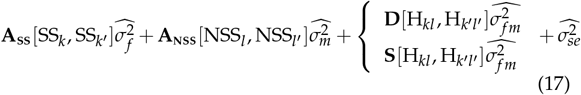

where the third component was modeled using both the **D** and **S** matrices, resulting in variants 4a and 4b. Note that the variance components 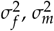, and 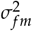 should be interpreted in relation to the covariance matrix used.

#### Assessment of predictive abilities

The predictive ability of each method was calculated using leave-one-out cross-validation, in which each single-cross hybrid was sequentially removed from the data set to serve as the validation set, while the remaining data were used to train the model and predict the excluded hybrid (James *et al*. 2013). To evaluate the robustness of the predictive methods under varying levels of parental information, we tested four training set configurations: (i) T2, where both parents of the excluded hybrid were present in the training set through other hybrids; (ii) T1F, where all hybrids sharing the same seed parent (SS) were excluded; (iii) T1M, where all hybrids sharing the same pollen parent (NSS) were excluded; and (iv) T0, where all hybrids involving either parent of the validation hybrid were excluded, ensuring no direct parental information was available in the training set (Technow *et al*. 2014; Kadam *et al*. 2016). To avoid confounding effects due to variation in training set size, we determined the largest training set size common to all four prediction configurations, which was 125 hybrids. Following this criterion, we held out one hybrid as the validation unit and randomly sampled 125 hybrids without replacement from the remaining 161 to form the training set. This procedure was repeated 30 times to ensure adequate resampling. Finally, we calculated Pearson’s correlation between the E-BLUEs of the hybrids obtained from Equation 4 and the genomic predicted genotypic values to assess the predictive ability of the methods.

### Model implementation

All linear models described in this study were fitted using the ASReml-R package v4 (The VSNi Team 2023). This package was particularly well-suited to our modeling framework because it allowed for the specification of complex variance-covariance structures for multi-level random effects. It also supported user-defined relationship matrices and facilitated the estimation of variance components using restricted maximum likelihood (REML). Moreover, ASReml-R provided AIC values for model comparison and enabled LRT to assess the significance of individual variance components.

## Results

### The optimal variance–covariance structure was trait-dependent

The comparison of GBLUP-based multi-kernel models, summarized in Table 2, revealed clear differences in model fit across traits (Table 4). Among the models fitted with the non-additive genomic relationship matrix (**D**), the one assuming environment-specific variance only for the three-way interaction term (M2-D) was favored for GY, EH, and SI. For PH, M1-D, which assumed homogeneous variances across environments, provided the best fit. In the case of AN, the preferred model was M3-D, which incorporated environment-specific variances for all interactions involving environments. Similarly, for the models based on the **S** matrix derived from additive genetic relationships between parental lines, M1-S yielded the lowest AIC for PH, whereas M2-S was superior for GY, EH, and SI, and M3-S was selected for AN.

**Table 4.**
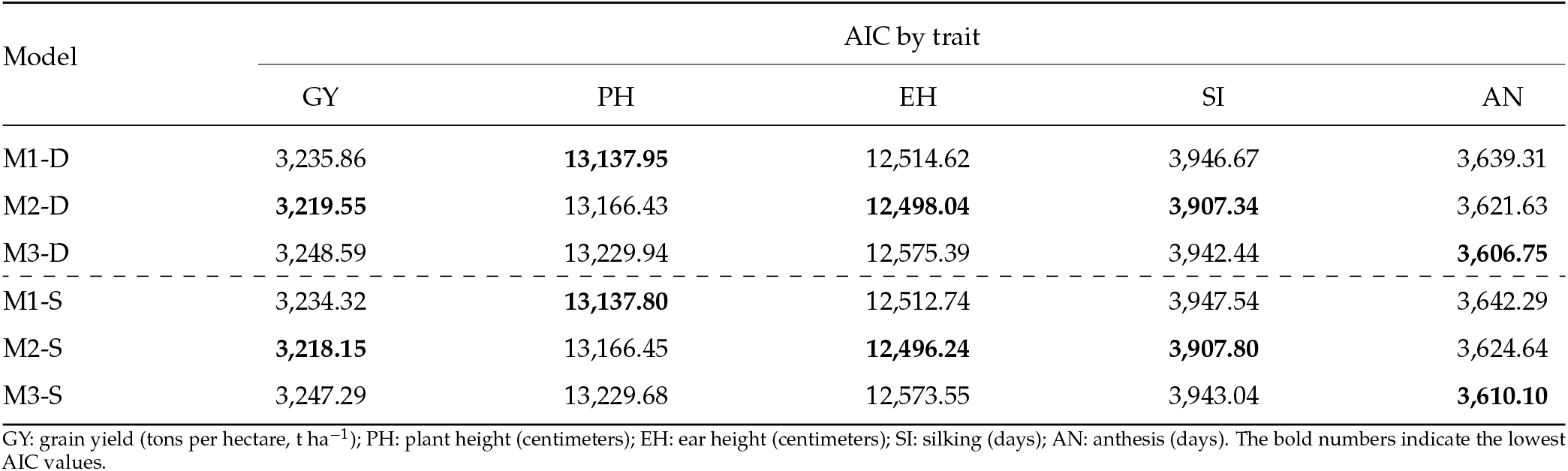
Akaike Information Criterion (AIC) values of GBLUP-based multi-kernel models by trait.

These findings show that the ideal variance-covariance structure depends on the trait being examined. A homogeneous variance is adequate for PH, while heterogeneous structures are more suitable for GY, EH, SI, and AN. Additionally, the similar AIC values and consistent rankings between the **D** and **S** matrices suggest that, although they are constructed differently, both parametrizations effectively capture the same underlying genetic architecture of the traits.

### GCA variance drives genetic variation in both groups, with some notable exceptions

Dissecting the total phenotypic variance when all lines were analyzed together highlighted significant variability in both heterotic groups, confirming that SS and NSS lines harbor GCA variance across all traits (Table 5). Both parametrizations used to estimate non-additive components, the **D** and **S** matrices, suggested the same underlying genetic architecture. The contribution of NSS lines to GCA variance was consistently greater than that of SS lines, particularly for PH and EH, where NSS explained nearly three times more variance. In contrast, the variance related to SCA was smaller than the GCA components, indicating that non-additive effects played a more limited role across the majority of traits under study. The main difference between the two parametrizations was in magnitude: the **S** matrix produced smaller estimates of SCA, reflecting a more conservative quantification of non-additive effects, without altering the overall conclusions.

**Table 5.** Estimates of variance components associated with general combining ability (GCA) and specific combining ability (SCA) obtained from the best-fitting GBLUP-based multi-kernel models for grain yield (GY, t^2^ ha^−2^), plant height (PH, cm^2^), ear height (EH, cm^2^), silking (SI, days^2^), and anthesis (AN, days^2^).

| Class<sup>1</sup> | Effect | Estimated variance components by trait |  |  |  |  |
| --- | --- | --- | --- | --- | --- | --- |
|  |  | GY | PH | EH | SI | AN |
| D | SS | 0.41 | 40.07 | 21.81 | 7.12 | 5.73 |
|  | NSS | 0.92 | 135.22 | 59.87 | 10.80 | 9.98 |
|  | SS × NSS | 0.31 | 14.57 | 5.82 | 0.47 | 0.57 |
| S | SS | 0.46 | 41.92 | 21.89 | 7.14 | 5.67 |
|  | NSS | 0.88 | 130.84 | 58.31 | 10.85 | 10.04 |
|  | SS × NSS | 0.07 | 3.28 | 1.30 | 0.11 | 0.13 |

SS: GCA of Stiff Stalk (SS) lines used as seed parents, reflecting additive effects; NSS: GCA of Non-Stiff Stalk (NSS) lines used as pollen parents, reflecting additive effects; and SS × NSS: SCA of single-cross hybrids between SS and NSS lines, reflecting non-additive effects.
<sup>1</sup> This column indicates whether variance components were estimated from models specified with the **D** matrix (Class D) or with the **S** matrix (Class S).

When examining the variance components within each maturity group (Table 6), the overall pattern observed in the combined data set was mostly maintained, with a few noteworthy differences among the groups. In the early maturity set, the SCA component for EH was not significant. In the intermediate group, the most important maturity group for US maize production, a critical result emerged: the SS component for GY, the most economically relevant trait, was not significant. In this case, the variance from SCA was greater than that from SS, indicating that SS contributed less to the total phenotypic variance compared to other sources of variation in the intermediate group. For the SI and AN, the variances of the SS and NSS intermediate lines were quite similar in magnitude, and SCA effects remained small. In the EH case, we found that the NSS variance was significant under the **D** model (59.34) but not under the **S** model (28.56). In the late maturity set, variance components were generally smaller than in the other groups, with some SCA estimates approaching zero. Nevertheless, several of these components were statistically significant according to the LRT, indicating that even minimal non-additive effects were detectable. For further details on the standard errors of the estimates and the LRT statistics from the model comparisons, see Tables S7 to S10.

**Table 6.** Estimates of variance components associated with general combining ability (GCA) and specific combining ability (SCA) obtained from the best-fitting GBLUP-based multi-kernel models within each maturity group for grain yield (GY, (t ha^−1^)^2^), plant height (PH, cm^2^), ear height (EH, cm^2^), silking (SI, days^2^), and anthesis (AN, days^2^).

| Maturity group | Class<sup>1</sup> | Effect | GY | PH | EH | SI | AN |
| --- | --- | --- | --- | --- | --- | --- | --- |
| Early | D | SS | 0.82 | 76.58 | 22.32 | 2.60 | 2.49 |
|  |  | NSS | 1.49 | 106.28 | 49.76 | 6.35 | 5.56 |
|  |  | SS × NSS | 0.18 | 2.20 | 0.71<sup>ns</sup> | 1.11 | 1.29 |
|  | S | SS | 0.76 | 79.46 | 23.05 | 2.93 | 2.83 |
|  |  | NSS | 1.42 | 107.63 | 50.74 | 6.25 | 5.37 |
|  |  | SS × NSS | 0.06 | 0.77 | 0.13<sup>ns</sup> | 0.34 | 0.40 |
| Intermediate | D | SS | 0.02<sup>ns</sup> | 7.19 | 23.16 | 3.92 | 2.97 |
|  |  | NSS | 0.71 | 145.74 | 59.34 | 2.58 | 2.41 |
|  |  | SS × NSS | 0.21 | 14.86 | 6.83 | 0.07 | 0.09 |
|  | S | SS | 0.03<sup>ns</sup> | 6.73 | 21.09 | 3.89 | 2.85 |
|  |  | NSS | 0.30 | 113.37 | 28.56<sup>ns</sup> | 2.22 | 2.05 |
|  |  | SS × NSS | 0.15 | 8.06 | 4.93 | 0.06 | 0.07 |
| Late | D | SS | 0.11 | 7.90 | 3.39 | 0.75 | 0.75 |
|  |  | NSS | 0.32 | 50.62 | 14.37 | 2.95 | 3.28 |
|  |  | SS × NSS | 0.01 | 0.42 | 0.29 | 0.06 | 0.08 |
|  | S | SS | 0.11 | 7.89 | 2.94 | 0.80 | 0.80 |
|  |  | NSS | 0.31 | 50.89 | 13.88 | 2.69 | 3.05 |
|  |  | SS × NSS | 0.01 | 0.40 | 0.39 | 0.05 | 0.06 |

SS: GCA of Stiff Stalk (SS) lines used as seed parents, reflecting additive effects; NSS: GCA of Non-Stiff Stalk (NSS) lines used as pollen parents, reflecting additive effects; SS × NSS: SCA of single-cross hybrids between SS and NSS lines, reflecting non-additive effects.
<sup>1</sup> This column indicates whether variance components were estimated from models specified with the **D** matrix (Class D) or with the **S** matrix (Class S).
<sup>ns</sup> Non-significant likelihood ratio test ( $p \geq 0.05$ ).

Figure 1 expands the variance partitioning by including the GEI terms, offering a more comprehensive view of the multiple sources of variation in GY. It should be noted that the Figure 1 does not include residual variance, as it was not re-estimated in model 9 but rather predefined using weights derived from single-trial analyses. Consequently, the partitions presented here only reflect the genetic and interaction sources of variation, and the “total variance” need be understood as the sum of SS, NSS, SS × NSS, SS × ENV, NSS × ENV, and SS × NSS × ENV components. In the combined data set, which does not separate maturity groups, the GEI effects accounted for a non-negligible proportion of the explained variance across models specified with either the **D** or the **S** formulation. GEI represented about 25% of the explained variance, while the SS effect explained roughly 10% and the SS × NSS interaction about 15%. These results indicate that environmental responsiveness was a significant driver of yield performance, in addition to the GCA variance contributed by the NSS lines.

**Figure 1.**
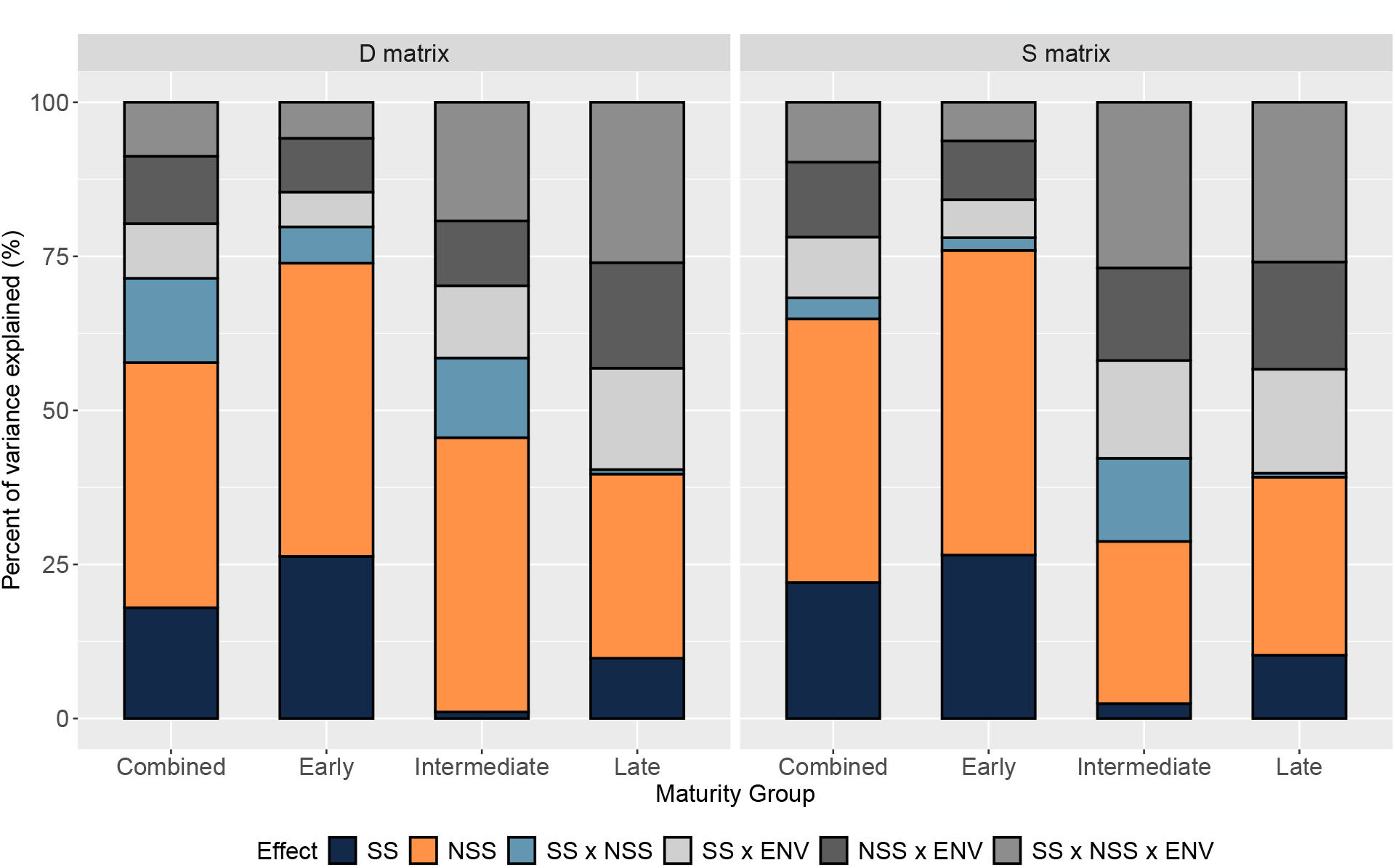
Percent of variance explained for grain yield (GY, tons per hectare, t ha^−1^). Variance components were estimated from models specified with either the **D** matrix or the **S** matrix. SS: general combining ability (GCA) of Stiff Stalk (SS) lines used as seed parents, reflecting additive effects; NSS: GCA of Non-Stiff Stalk (NSS) lines used as pollen parents, reflecting additive effects; SS × NSS: specific combining ability (SCA) of single-cross hybrids between SS and NSS lines, reflecting non-additive effects; SS × ENV: interaction between SS lines and environments; NSS × ENV: interaction between NSS lines and environments; SS × NSS × ENV: three-way interaction among SS, NSS, and environments. Results are shown for all hybrids (Combined) as well as separately by maturity group (Early, Intermediate, Late).

By maturity groups, notable shifts in the proportion of variance explained by GEI became evident (Figure 1). In early hybrids, nearly 70% of the observed variance was attributed to SS and NSS lines, while GEI accounted for only a minor proportion, indicating that hybrid performance across environments was relatively stable. For intermediate hybrids, the proportion of variance explained by GEI increased compared with the early group. Notably, when considering all three interaction sources, GEI accounted for more than 40% of the total variance. The highest percentage of GEI was found in the late group, where interaction terms represented more than 50% of the total variance. The increase can be largely attributed to the wider environmental range. In our analysis of the raw dataset, we found that late hybrids were grown not just in environments classified as late, but also in those categorized as early and intermediate. Such cross-testing likely accentuated differences in hybrid adaptation and magnified GEI signals.

Similar variance partitioning plots for the other traits are provided in Supplementary Figures S3, S4, S5, and S6. Overall, the patterns were comparable to those described for GY, although trait-specific differences in GEI magnitude were observed. For traits modeled with heterogeneous variance structures for the interaction terms (see Table 2), we calculated a single variance estimate using weighted averages. In this process, the variance estimated for each environment was multiplied by the number of observations available in that environment. The resulting products were then summed across all environments. This approach ensured that the resulting estimates were directly comparable to those for traits, such as PH, which were modeled under homogeneous variance assumptions.

### Predictive ability hinges on parental information and the method used, with GCA effects providing sufficient accuracy in this study

Across traits, predictive ability declined as parental information was progressively removed from the training data (Figure 2). Configuration T2, where training and validation sets shared the strongest connectedness, produced the highest correlations between E-BLUEs and genomic predicted genotypic values. In contrast, removing seed parents (SS) in T1F or pollen parents (NSS) in T1M resulted in a moderate decline in performance. The T0 configuration generally produced the lowest correlations, although its impact varied across traits and methods, with some combinations maintaining comparable levels to T1F and T1M. Across all traits, GY demonstrated the lowest predictive abilities across configurations and was most significantly affected by the absence of parental information. Meanwhile, in line with their greater heritability (Supplementary Table S5), the traits related to flowering time (SI and AN) exhibited the highest predictive abilities, whereas PH and EH had predictive abilities that intermediate relative to the other traits.

**Figure 2.**
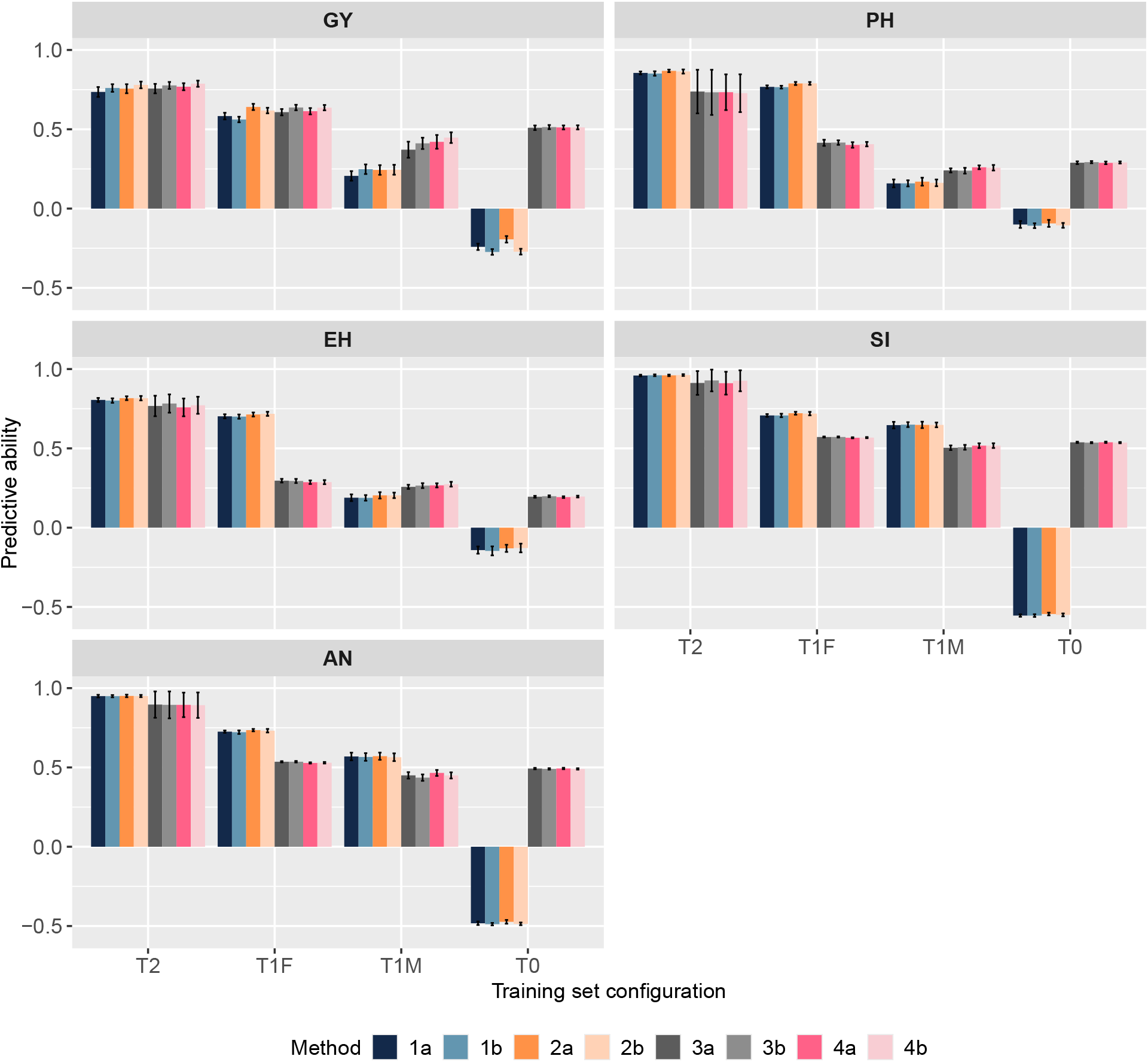
Predictive ability for untested single-cross hybrids in grain yield (GY), plant height (PH), ear height (EH), silking (SI), and anthesis (AN), assessed under four training-set configurations: T2 (both parents of the validation hybrid represented via other crosses), T1F (all hybrids sharing the same seed parent excluded), T1M (all hybrids sharing the same pollen parent excluded), and T0 (no direct parental information in the training set). Methods 1–2 correspond to GBLUP-based multi-kernel models, where method 1 includes only general combining ability (GCA) and method 2 consists of both GCA and specific combining ability (SCA). Methods 3–4 are based on the covariance between tested and untested single-cross hybrids, considering either the additive relationship matrix only (method 3) or both additive and non-additive relationship matrices (method 4). Results are shown for models fitted using either the **D** matrix (a) or the **S** matrix (b). Additional model details are provided in Table 3.

In comparing the methods that utilize GBLUP-based multi-kernel models (1a, 1b, 2a, and 2b) with those that focus on the covariance between tested and untested single-cross hybrids (3a, 3b, 4a, and 4b), clear differences emerged across traits and training configurations (Figure 2). Under T2, methods using GBLUP-based multi-kernel models consistently outperformed covariance-based methods for most traits. The only exception to this trend was GY, where the predictive abilities of the two strategies were nearly overlapping. The T1F training set configuration followed the same general pattern as T2, but the advantage of GBLUP-based multi-kernel models became more pronounced, particularly for PH and EH, where the gap between the two approaches was larger. Results under T1M were more trait-dependent: GBLUP-based multi-kernel models showed superior performance for SI and AN, while covariance-based models achieved higher predictive abilities for PH, EH, and GY. The most contrasting outcomes were observed in the T0 scenario, which was the strictest. In this case, GBLUP-based multi-kernel models often yielded negative accuracy values across traits, particularly for GY. In turn, the covariance-based methods continued to show higher predictive abilities for SI, AN, and GY, with moderate performance for PH and EH. This comparison clearly shows that GBLUP-based multi-kernel models generally perform better when parental information is available, particularly for traits with higher heritability, such as SI and AN. Conversely, covariance-based predictions tend to be more stable than GBLUP-based multi-kernel models in situations where direct parental representation is lacking.

The addition of SCA effects (2a–2b) or non-additive matrices (4a–4b) produced negligible differences in predictive ability, indicating that non-additive effects played a limited role in this study (Figure 2). These outcomes are consistent with the variance component estimates from the multi-kernel models (Table 5), where GCA effects explained the largest portion of genetic variance across traits, particularly for PH, EH, SI, and AN in both SS and NSS heterotic groups. The predominance of GCA variance in these populations provides a natural rationale for the strong performance of models that rely on GCA to predict hybrid performance. Moreover, predictive abilities were nearly identical between variants that used the **D** matrix (a) and those that used the **S** matrix (b), also suggesting that these matrices capture the same underlying genetic architecture. This similarity was anticipated, as both parameterizations provided nearly identical variance components in most instances (Tables 5 and 6).

## Discussion

The reduction of variance components associated with GCA within heterotic groups and the challenge of predicting untested single crosses at early stages remain central issues in maize breeding programs. Using GBLUP-based multi-kernel models, this study examined how GCA variance is distributed within SS and NSS germplasm and how parental information shapes predictive performance. The findings indicate that GCA remains a significant source of genetic signal for most traits. However, the study also highlights context-dependent limitations, particularly for grain yield in SS germplasm, and shows that predictive ability is strongly influenced by the degree of parental connectivity.

### Maintaining variance associated with GCA effects within heterotic groups is critical for sustained hybrid breeding progress

Additive variance refers to the portion of total genetic variance attributable to the average effects of alleles (Falconer 1996). It represents the effects that can be consistently transmitted from parents to progeny, and therefore it underlies the predictable response to selection (Lynch et al. 1998). In the classic breeder’s equation, the expected genetic gain is directly proportional to the amount of additive variance present in the population (Bernardo et al. 2002). For this reason, the long-term success of maize hybrid breeding programs depends on maintaining additive variance within heterotic groups (Melchinger 1999; Hallauer et al. 2010). If this source of variance becomes depleted, further genetic gain cannot be achieved (Walsh and Lynch 2018).

The GCA effects of parental lines capture differences in their average performance across multiple crosses, and these differences arise primarily from additive gene action. Based on this relationship, the magnitude of the GCA variance provides an indirect yet informative measure of the additive-related genetic variation available within each heterotic group. Thus, monitoring changes in GCA variance offers practical insight into the status and long-term sustainability of additive variance within breeding groups (Orton 2020).

The set of SS and NSS inbred lines analyzed in this study represents elite parental germplasm developed and released for hybrid breeding between the late 1980s and mid-1990s (Beckett et al. 2017). Over the decades that followed, repeated cycles of selection and the recycling of inbred lines within these heterotic groups have led to significant genetic progress. One clear example is the improvement in grain yield under drought-stress conditions (Gaffney *et al*. 2015). Recent analyses conducted in the US Corn Belt have shown that maize yields have increased in both favorable and stressed environments, averaging an annual gain of approximately 1.2%. Also, newer hybrids demonstrated improved drought resistance during the grain-filling period (Zhao et al. 2025). Nevertheless, this strategy of repeatedly drawing upon a relatively closed pool of parental lines, combined with high selection intensities, increases the risk of allele fixation, reduces opportunities to discover new favorable combinations, and can lead to a gradual erosion of genetic diversity (Hallauer 1992).

Our findings demonstrate that, for the majority of traits evaluated, both SS and NSS heterotic groups have maintained significant GCA variance, whether analyzed jointly or within maturity groups. Similar findings were reported by Bornowski *et al*. (2021), who analyzed the SS germplasm using whole-genome assemblies and SNP-based diversity metrics, demonstrating that the pool retains significant genetic and genomic variation. Since their research focused on inbred lines, this residual diversity largely reflects the additive variance present within the population (Falconer 1996; Lynch et al. 1998) (Falconer 1996; Lynch et al. 1998). Our results, however, also highlight a critical concern: among the SS lines represented in our dataset, the GCA variance for GY in the intermediate maturity group was not statistically significant. This maturity group is crucial for the US Corn Belt, where intermediate hybrids dominate commercial fields and account for nearly one-third of global maize production (Zhao *et al*. 2025). Moreover, GY is the most important trait for the commercial success of maize. The lack of GCA variance for GY in intermediate lines indicates a significant weakness in the SS germplasm, particularly in light of the increasing climatic challenges projected for the US Corn Belt (Zhao *et al*. 2025).

Looking forward, the concern extends beyond the intermediate set. The GCA variance still present in SS and NSS germplasm for other traits and maturity groups is a finite resource that, as discussed above, may gradually diminish over time (Hallauer 1992; Goodman 2005). To safeguard long-term progress, it is therefore essential not only to exploit the remaining variance but also to adopt strategies that prevent its depletion (Goodman 1999; Swarup *et al*. 2021; Salgotra and Chauhan 2023). Modern breeding tools, such as genomic selection, play an important role in maintaining genetic diversity by limiting allele fixation and controlling relatedness within recurrent breeding programs (Li *et al*. 2022). Complementary to these approaches, pre-breeding efforts, like the Germplasm Enhancement of Maize project, have been instrumental in broadening the genetic base of commercial maize in the US by incorporating novel and useful germplasm collected worldwide (Pollak and Salhuana 2000).

### The transient role of non-additive variance captured by SCA in long-term genetic gain and the pervasive influence of GEI

Shull (1908) and East (1908) independently demonstrated that inbred maize lines exhibited reduced vigor and productivity, whereas crosses between distinct inbreds produced offspring with markedly improved performance due to a phenomenon later termed heterosis. Since then, unraveling the genetic basis of heterosis has been a major focus in maize research, with dominance, overdominance, and epistasis recognized as key mechanisms contributing to the superior performance of hybrids. In maize, these non-additive mechanisms are not mutually exclusive and are captured by SCA effects (Garcia *et al*. 2025).

The magnitude of the SCA effects determines how much hybrid performance deviates from what is expected based on parental GCA effects alone (Hallauer 1992; Hallauer *et al*. 2010). In our study, the variance attributed to SCA was consistently smaller than the variance explained by GCA across traits. This finding aligns with the theoretical expectations of Reif *et al*. (2007), who showed that the ratio of dominance to additive variance decreases with increasing interpopulation divergence, resulting in a greater prevalence of GCA over SCA in divergent heterotic groups. A simulation study also demonstrated that both GCA and SCA variances tend to decrease across recurrent cycles of selection due to the strong selection intensity, with the reduction being particularly pronounced for SCA (Melchinger and Frisch 2023). These theoretical insights provide a compelling explanation for the lower magnitude of SCA variance observed in our data set and highlight that, while SCA contributes to the expression of heterosis, its role is transient compared with the more enduring additive effects in long-term genetic gain. Moreover, Melchinger and Frisch (2023) emphasized that the relative importance of SCA varies across traits, and our results are in line with this expectation: we found a higher SCA contribution for GY than for flowering-related traits, confirming that the expression of non-additive variance is largely trait-dependent.

The GEI component represents a fundamental source of variation in hybrid breeding because it captures the differential response of genotypes across contrasting environments. Its presence complicates selection decisions, as superior performance in one environment does not necessarily translate into broad adaptation. Still, it also offers opportunities to exploit specific adaptation when target populations of environments are clearly defined (Van Eeuwijk et al. 2016). Therefore, accounting for GEI, whether through explicit interaction terms or implicit variance-covariance structures, is essential for accurate estimation of genetic effects and reliable prediction of hybrid performance. Numerous studies across maize have demonstrated that modeling GEI improves predictive ability and provides more realistic assessments of adaptation and stability, particularly for complex traits such as grain yield (Zhang et al. 2015; Dias et al. 2018).

The GEI component was modeled explicitly in this study, which allowed us to directly quantify its contribution to the total genetic variance. This approach revealed clear differences across maturity groups, with the most pronounced GEI detected in late hybrids. A likely explanation is that these environments included not only late-maturity hybrids but also early and intermediate ones, thereby amplifying differential responses and magnifying interaction effects. Historically, maturity groups have been central to US maize breeding because they align hybrids with the latitudinal gradient of production environments, ensuring that flowering and grain-filling occur under favorable seasonal conditions (Massigoge *et al*. 2023). The stronger GEI observed in late hybrids thus reflects both the broader environmental range in which they were tested and the intrinsic importance of maturity groups in structuring hybrid adaptation. These outcomes emphasize that explicitly accounting for GEI provides a more accurate understanding of the distribution of genetic variance and highlights its decisive role in shaping hybrid performance across environments.

### Integrating mating designs and genomic relationship matrices for the orthogonal partition of genetic variance

In maize, dominance significantly influences hybrid performance, so modeling both additive and non-additive effects is crucial for understanding trait architecture and guiding breeding strategies (Nazarian and Gezan 2016; Dias et al. 2018). One effective approach to achieve this is through suitable mating designs, such as the North Carolina design II, also known as a factorial design (Comstock and Robinson 1948). In this design, a set of pollen parents is systematically crossed with multiple seed parents, resulting in progenies that represent two distinct levels of relatedness: half-sib families, which come from either the pollen or seed parent, and full-sib families within each specific pollen-seed combination. Because both relationships are present, breeders can obtain complementary estimates of additive variance attributable to the parental lines and non-additive variance resulting from specific parental combinations (Bernardo et al. 2002).

In scenarios where orthogonality is not ensured by design, molecular marker–based relationship matrices provide breeders with a robust framework for estimating genetic effects and their associated variance components (Nazarian and Gezan 2016; Bouvet et al. 2016; Dias et al. 2018). The additive genomic relationship matrix quantifies the proportion of alleles shared between individuals at each locus, whereas the dominance genomic relationship matrix reflects covariances due to specific allelic combinations at that locus. Although several methodologies can be used to generate these matrices, the most widely adopted are those proposed by VanRaden (2008) for additive effects and by Vitezica *et al*. (2013) and Muñoz *et al*. (2014) for dominance effects. In these approaches, the matrices are explicitly centered and parameterized with respect to allele frequencies, ensuring that the expected covariance between additive and dominance components is zero. Consequently, the two sources of variation are statistically independent, enabling breeders to obtain orthogonal estimates of additive and dominance effects and, ultimately, a clear partitioning of the underlying genetic variance.

We were able to partition genetic variance into components associated with GCA effects and non-additive effects because the single-cross hybrids used in this study were generated in a factorial design. Also, incorporating additive and non-additive matrices into the model further reinforced this orthogonality. Beyond ensuring independence among sources of variation, these matrices provided a practical framework for anticipating the performance of untested hybrids, an outcome central to the efficiency and scalability of modern maize breeding. Regarding the modeling of non-additive components, both **D** and **S** produced nearly identical patterns for all outcomes. This result suggests both matrices captured the same overall trends in underlying trait genetic architecture. The main difference lies in the magnitude of the variance components, which can be attributed to the mathematical properties of the two parameterizations: whereas the **D** matrix directly models non-additive deviations based on allele frequencies, the **S** matrix rescales these effects through a transformation of the additive relationships, typically yielding more conservative estimates of non-additive variance.

### Parental information and GCA variance are decisive for maize genomic prediction

The existence of a direct genetic relationship between the training and validation populations is fundamental to the success of genomic selection (Desta and Ortiz 2014; Ferrão *et al*. 2017), and this became evident in our study. As parental information was progressively removed from the training set, predictive ability declined, underscoring the central role of genetic connectedness in capturing the additive signal required for accurate predictions. This pattern, reflected in the superior performance of T2 over T1F, T1M, and especially T0, is consistent with previous research on maize hybrid prediction (Technow *et al*. 2014; Kadam *et al*. 2016). Using maize data–based simulations, Melchinger and Frisch (2023) also showed that prediction accuracy is maximized when more parents are represented in the training population, even if each contributes only a single cross (Technow *et al*. 2014; Kadam *et al*. 2016). Another factor affecting genomic selection accuracy is the amount of additive genetic variance (Desta and Ortiz 2014; Ferrão *et al*. 2017), which was highlighted by the sharper decline under T1M. In this scenario, the exclusion of NSS parents, who contributed most to this variance, markedly reduced predictive ability when compared to T1F.

GBLUP-based models (1a, 1b, 2a, and 2b) generally outperformed the covariance approaches (3a, 3b, 4a, and 4b), confirming their effectiveness when parental information was available. Under T0, however, they produced negative correlations for all traits, whereas the covariance approaches still yielded positive values. This discrepancy can be attributed to differences in how the two methods estimate effects. In GBLUP-based models, the GCA effects of individual parents and the SCA effects of their combinations are estimated using mixed-model equations (Henderson 1975). When neither parent of a hybrid is represented in the training set (T0), the model lacks anchors to predict these effects, which are consequently shrunk toward zero (Piepho *et al*. 2008). Shrinkage of the estimates, combined with allele-frequency centering of genomic matrices, can lead to negative predictive ability, even when genomic relationship matrices retain residual similarity. In contrast, covariance-based approaches generate predictions by projecting the phenotypes of tested hybrids onto those of untested hybrids using covariance blocks 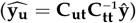. In this framework, even if the exact parents of a hybrid are not represented in the training set, genomic similarity with other related parents or hybrids ensures non-zero covariance values. As a result, this information is directly incorporated into the predictions, resulting in stable, positive accuracies even in the absence of explicit parental overlap (Bernardo 1996).

Kadam *et al*. (2016) also evaluated genomic prediction under T0 conditions but reported positive accuracies. This difference can be attributed to population structure: their study involved 312 hybrids derived from six biparental families, which created a highly interconnected dataset. Even when the exact parents of a hybrid were excluded, other hybrids sharing closely related parents were still represented, sustaining non-zero covariance and positive predictive ability. In our case, the diverse SS and NSS parental lines produced a population with much lower indirect connectivity, making our T0 scenario more stringent.

Trait-specific differences in predictive ability reflected their underlying genetic architecture. Plant architectural and flowering traits such as PH, EH, SI, and AN showed higher accuracies because they are largely governed by additive genetic variance and exhibit higher heritability, allowing models to capture most of the genetic signal from parental information. By contrast, GY showed lower predictive ability and was disproportionately affected by the removal of parental connectivity, consistent with its smaller additive component and greater sensitivity to GEI and residual variance (Bernardo et al. 2002). These results highlight the importance of modeling GEI in genomic selection frameworks for complex traits such as GY, as failure to account for this source of variation can undermine prediction accuracy (Dias et al. 2018).

The limited gains from including non-additive effects in our models should not be interpreted as evidence that non-additive effect is universally negligible. In quantitative genetics, the relevance of additive and non-additive components always depends on the population and context in which they are defined. For this germplasm, GCA variance clearly accounted for most of the observed phenotypic differences, which explains why models built only on GCA performed so well. However, other studies have shown meaningful increases in predictive ability when non-additive effects were incorporated (Dias et al. 2018). This underscores the need for breeders to evaluate the genetic architecture of their target populations before deciding whether to include such terms, as the contribution of non-additive effect is not a universal constant but a property of the specific genetic material under study (Garcia et al. 2025).

## Conclusion

The GBLUP-based multi-kernel models developed in this study revealed that the SS and NSS parental lines represented in our dataset possess significant GCA variance, with NSS contributing more across various traits. However, in the most commercially relevant maturity group in the US Corn Belt, the intermediate SS lines, showed no significant GCA variance for grain yield. This putatively small amount of GCA variance limits the potential for sustained genetic improvement and underscores the need to expand the genetic base of SS germplasm to ensure long-term breeding progress. Furthermore, the GBLUP-based multi-kernel models performed well in T2, T1F, and T1M, where at least one parent was included in the training set, but poorly in T0, where no parental information was available. Since breeding programs typically operate under scenarios more comparable to T2, T1F, and T1M, our results demonstrate the practical value of this framework for improving the efficiency of maize hybrid breeding.

## Code and data availability

All scripts and processed data supporting this study are publicly available via Zenodo at https://doi.org/10.5281/zenodo.17917188. The Zenodo record is linked to the corresponding GitHub repository, which includes documentation and instructions to retrieve the raw phenotypic and genotypic data from the G2F Data Commons (CyVerse).

## Authors contributions

JCGS developed and conducted the data analysis and prediction pipeline and wrote the manuscript. MB, MM, JE, EL, and AL conceived and designed the study. SF contributed expertise to data analysis and interpretation. MB, JE, and EL produced the seed, and MB, JE, EL, and CH conducted parts of the field experiments. All authors reviewed and edited the manuscript and approved the final version for publication.

## Acknowledgements

The authors used Grammarly software to assist with grammar and language editing during manuscript preparation. The authors reviewed all edits and take full responsibility for the content of the manuscript.

