## Supplemental Materials for "Dissecting genetic variance structure and evaluating genomic prediction models for single-cross hybrids derived from Stiff Stalk and Non-Stiff Stalk maize heterotic groups"

### Supplementary Information

**Table S1**: List of single-cross hybrids derived from Stiff Stalk (SS) × Non-Stiff Stalk (NSS) inbred lines that were exclusively assigned to the early maturity group.

| SS | NSS | Single Cross | Maturity Group |
|---|---|---|---|
| LH145 | PHR25 | LH145/PHR25 | Early |
| CG120 | PHR25 | CG120/PHR25 | Early |
| PHRE1 | PHR25 | PHRE1/PHR25 | Early |
| PHB47 | PHR25 | PHB47/PHR25 | Early |
| PHHV4 | PHR25 | PHHV4/PHR25 | Early |
| S8324 | PHR25 | S8324/PHR25 | Early |
| CG120 | PHZ51 | CG120/PHZ51 | Early |
| PHHV4 | PHZ51 | PHHV4/PHZ51 | Early |
| S8324 | PHZ51 | S8324/PHZ51 | Early |
| CG120 | LH185 | CG120/LH185 | Early |
| PHHV4 | LH185 | PHHV4/LH185 | Early |
| PHHV4 | PHK56 | PHHV4/PHK56 | Early |
| S8324 | PHK56 | S8324/PHK56 | Early |
| LH145 | LH162 | LH145/LH162 | Early |
| CG120 | LH162 | CG120/LH162 | Early |
| PHRE1 | LH162 | PHRE1/LH162 | Early |
| PHB47 | LH162 | PHB47/LH162 | Early |
| PHHV4 | LH162 | PHHV4/LH162 | Early |
| S8324 | LH162 | S8324/LH162 | Early |
| LH145 | CG123 | LH145/CG123 | Early |
| CG120 | CG123 | CG120/CG123 | Early |
| PHRE1 | CG123 | PHRE1/CG123 | Early |
| PHB47 | CG123 | PHB47/CG123 | Early |
| PHHV4 | CG123 | PHHV4/CG123 | Early |
| S8324 | CG123 | S8324/CG123 | Early |
| CG120 | PHW03 | CG120/PHW03 | Early |
| PHB47 | PHW03 | PHB47/PHW03 | Early |
| PHHV4 | PHW03 | PHHV4/PHW03 | Early |
| LH145 | PHK05 | LH145/PHK05 | Early |
| CG120 | PHK05 | CG120/PHK05 | Early |
| PHB47 | PHK05 | PHB47/PHK05 | Early |
| PHHV4 | PHK05 | PHHV4/PHK05 | Early |
| S8324 | PHK05 | S8324/PHK05 | Early |

**Table S2**: List of single-cross hybrids derived from Stiff Stalk (SS) × Non-Stiff Stalk (NSS) inbred lines that were exclusively assigned to the intermediate maturity group.

| SS | NSS | Single Cross | Maturity Group |
|---|---|---|---|
| PHN66 | PHG83 | PHN66/PHG83 | Intermediate |
| PHB47 | PHG83 | PHB47/PHG83 | Intermediate |
| PHW52 | PHG83 | PHW52/PHG83 | Intermediate |
| LH198 | PHG83 | LH198/PHG83 | Intermediate |
| LH195 | PHG83 | LH195/PHG83 | Intermediate |
| PHN66 | Q381 | PHN66/Q381 | Intermediate |
| PHB47 | Q381 | PHB47/Q381 | Intermediate |
| LH198 | Q381 | LH198/Q381 | Intermediate |
| PHN66 | PHN82 | PHN66/PHN82 | Intermediate |
| LH198 | PHN82 | LH198/PHN82 | Intermediate |
| PHN66 | PHG29 | PHN66/PHG29 | Intermediate |
| PHW52 | PHG29 | PHW52/PHG29 | Intermediate |
| LH198 | PHG29 | LH198/PHG29 | Intermediate |
| LH195 | PHG29 | LH195/PHG29 | Intermediate |
| PHN66 | LH82 | PHN66/LH82 | Intermediate |
| PHW52 | LH82 | PHW52/LH82 | Intermediate |
| LH198 | LH82 | LH198/LH82 | Intermediate |
| LH195 | LH82 | LH195/LH82 | Intermediate |
| PHN66 | PHZ51 | PHN66/PHZ51 | Intermediate |
| LH198 | PHZ51 | LH198/PHZ51 | Intermediate |
| PHN66 | PHR55 | PHN66/PHR55 | Intermediate |
| PHB47 | PHR55 | PHB47/PHR55 | Intermediate |
| LH198 | PHR55 | LH198/PHR55 | Intermediate |
| PHN66 | LH185 | PHN66/LH185 | Intermediate |
| LH198 | LH185 | LH198/LH185 | Intermediate |
| PHN66 | PHK76 | PHN66/PHK76 | Intermediate |
| PHB47 | PHK76 | PHB47/PHK76 | Intermediate |
| PHW52 | PHK76 | PHW52/PHK76 | Intermediate |
| LH198 | PHK76 | LH198/PHK76 | Intermediate |
| LH195 | PHK76 | LH195/PHK76 | Intermediate |
| PHN66 | LH38 | PHN66/LH38 | Intermediate |
| PHB47 | LH38 | PHB47/LH38 | Intermediate |
| PHW52 | LH38 | PHW52/LH38 | Intermediate |
| LH198 | LH38 | LH198/LH38 | Intermediate |
| LH195 | LH38 | LH195/LH38 | Intermediate |
| PHN66 | PHK56 | PHN66/PHK56 | Intermediate |
| PHW52 | PHK56 | PHW52/PHK56 | Intermediate |
| LH198 | PHK56 | LH198/PHK56 | Intermediate |
| LH195 | PHK56 | LH195/PHK56 | Intermediate |
| PHN66 | PHG47 | PHN66/PHG47 | Intermediate |
| PHW52 | PHG47 | PHW52/PHG47 | Intermediate |
| LH198 | PHG47 | LH198/PHG47 | Intermediate |
| LH195 | PHG47 | LH195/PHG47 | Intermediate |
| PHN66 | LH51 | PHN66/LH51 | Intermediate |
| PHB47 | LH51 | PHB47/LH51 | Intermediate |
| PHW52 | LH51 | PHW52/LH51 | Intermediate |
| LH198 | LH51 | LH198/LH51 | Intermediate |
| LH195 | LH51 | LH195/LH51 | Intermediate |

**Table S3**: List of single-cross hybrids derived from Stiff Stalk (SS) × Non-Stiff Stalk (NSS) inbred lines that were exclusively assigned to the late maturity group.

| SS | NSS | Single Cross | Maturity Group |
|---|---|---|---|
| PHHB9 | PHN47 | PHHB9/PHN47 | Late |
| PHV63 | PHN47 | PHV63/PHN47 | Late |
| PHP38 | PHN47 | PHP38/PHN47 | Late |
| LH195 | PHN47 | LH195/PHN47 | Late |
| PHW52 | PHN47 | PHW52/PHN47 | Late |
| PHHB9 | Q381 | PHHB9/Q381 | Late |
| PHV63 | Q381 | PHV63/Q381 | Late |
| PHP38 | Q381 | PHP38/Q381 | Late |
| PHHB9 | PHN82 | PHHB9/PHN82 | Late |
| PHV63 | PHN82 | PHV63/PHN82 | Late |
| PHP38 | PHN82 | PHP38/PHN82 | Late |
| PHHB9 | PHW30 | PHHB9/PHW30 | Late |
| PHV63 | PHW30 | PHV63/PHW30 | Late |
| PHP38 | PHW30 | PHP38/PHW30 | Late |
| PHHB9 | PHR63 | PHHB9/PHR63 | Late |
| PHV63 | PHR63 | PHV63/PHR63 | Late |
| PHP38 | PHR63 | PHP38/PHR63 | Late |
| LH195 | PHR63 | LH195/PHR63 | Late |
| PHW52 | PHR63 | PHW52/PHR63 | Late |
| PHHB9 | PHM49 | PHHB9/PHM49 | Late |
| PHV63 | PHM49 | PHV63/PHM49 | Late |
| PHP38 | PHM49 | PHP38/PHM49 | Late |
| LH195 | PHM49 | LH195/PHM49 | Late |
| PHW52 | PHM49 | PHW52/PHM49 | Late |
| PHHB9 | PHW53 | PHHB9/PHW53 | Late |
| PHV63 | PHW53 | PHV63/PHW53 | Late |
| PHP38 | PHW53 | PHP38/PHW53 | Late |
| LH195 | PHW53 | LH195/PHW53 | Late |
| PHW52 | PHW53 | PHW52/PHW53 | Late |
| PHHB9 | PHZ51 | PHHB9/PHZ51 | Late |
| PHV63 | PHZ51 | PHV63/PHZ51 | Late |
| PHHB9 | PHR55 | PHHB9/PHR55 | Late |
| PHV63 | PHR55 | PHV63/PHR55 | Late |
| PHP38 | PHR55 | PHP38/PHR55 | Late |
| PHHB9 | PHR03 | PHHB9/PHR03 | Late |
| PHV63 | PHR03 | PHV63/PHR03 | Late |
| PHP38 | PHR03 | PHP38/PHR03 | Late |
| LH195 | PHR03 | LH195/PHR03 | Late |
| PHW52 | PHR03 | PHW52/PHR03 | Late |
| PHHB9 | LH185 | PHHB9/LH185 | Late |
| PHV63 | LH185 | PHV63/LH185 | Late |
| PHP38 | LH185 | PHP38/LH185 | Late |
| PHHB9 | PHP60 | PHHB9/PHP60 | Late |
| PHV63 | PHP60 | PHV63/PHP60 | Late |
| PHP38 | PHP60 | PHP38/PHP60 | Late |
| LH195 | PHP60 | LH195/PHP60 | Late |
| PHW52 | PHP60 | PHW52/PHP60 | Late |
| PHHB9 | LH210 | PHHB9/LH210 | Late |
| PHV63 | LH210 | PHV63/LH210 | Late |
| PHP38 | LH210 | PHP38/LH210 | Late |
| LH195 | LH210 | LH195/LH210 | Late |
| PHW52 | LH210 | PHW52/LH210 | Late |
| PHHB9 | PHJ65 | PHHB9/PHJ65 | Late |
| PHV63 | PHJ65 | PHV63/PHJ65 | Late |
| PHP38 | PHJ65 | PHP38/PHJ65 | Late |
| LH195 | PHJ65 | LH195/PHJ65 | Late |
| PHW52 | PHJ65 | PHW52/PHJ65 | Late |
| PHHB9 | PHM57 | PHHB9/PHM57 | Late |
| PHV63 | PHM57 | PHV63/PHM57 | Late |
| PHP38 | PHM57 | PHP38/PHM57 | Late |
| LH195 | PHM57 | LH195/PHM57 | Late |
| PHW52 | PHM57 | PHW52/PHM57 | Late |

**Table S4**: List of single-cross hybrids derived from Stiff Stalk (SS) × Non-Stiff Stalk (NSS) lines that were simultaneously assigned to more than one maturity group.

| SS | NSS | Single Cross | Maturity Group |
|---|---|---|---|
| PHB47 | PHN82 | PHB47/PHN82 | Early/Intermediate |
| PHB47 | PHG29 | PHB47/PHG29 | Early/Intermediate |
| PHB47 | LH82 | PHB47/LH82 | Early/Intermediate |
| PHB47 | PHZ51 | PHB47/PHZ51 | Early/Intermediate |
| PHB47 | LH185 | PHB47/LH185 | Early/Intermediate |
| PHB47 | PHK56 | PHB47/PHK56 | Early/Intermediate |
| PHB47 | PHG47 | PHB47/PHG47 | Early/Intermediate |
| PHW52 | Q381 | PHW52/Q381 | Intermediate /Late |
| LH195 | Q381 | LH195/Q381 | Intermediate /Late |
| PHW52 | PHN82 | PHW52/PHN82 | Intermediate /Late |
| LH195 | PHN82 | LH195/PHN82 | Intermediate /Late |
| PHW52 | PHW30 | PHW52/PHW30 | Intermediate /Late |
| LH195 | PHW30 | LH195/PHW30 | Intermediate /Late |
| PHW52 | PHZ51 | PHW52/PHZ51 | Intermediate /Late |
| LH195 | PHZ51 | LH195/PHZ51 | Intermediate /Late |
| PHW52 | PHR55 | PHW52/PHR55 | Intermediate /Late |
| LH195 | PHR55 | LH195/PHR55 | Intermediate /Late |
| PHW52 | LH185 | PHW52/LH185 | Intermediate /Late |
| LH195 | LH185 | LH195/LH185 | Intermediate /Late |

**Table S5**: Broad-sense heritability ($H^2$) and coefficient of variation ($CV$, %) for grain yield (GY, t ha$^{-1}$), plant height (PH, centimeters), ear height (EH, centimeters), silking (SI, days), and anthesis (AN, days) across environments retained for analysis, i.e., those not excluded due to excessively high $CV$ (>25%) or near-zero $H^2$ values.

| Environment | GY | | PH | | EH | | SI | | AN | |
|---|---|---|---|---|---|---|---|---|---|---|
| | $H^2$ | $CV$ | $H^2$ | $CV$ | $H^2$ | $CV$ | $H^2$ | $CV$ | $H^2$ | $CV$ |
| 2016_ARH1 | 0.61 | 13.29 | 0.66 | 7.03 | 0.29 | 13.02 | 0.92 | 1.65 | 0.92 | 1.49 |
| 2016_ARH2 | 0.39 | 21.77 | 0.50 | 6.66 | 0.37 | 12.00 | 0.88 | 1.99 | 0.87 | 1.96 |
| 2016_DEH1 | 0.76 | 13.90 | 0.70 | 5.09 | 0.64 | 11.27 | 0.57 | 2.38 | 0.59 | 2.27 |
| 2016_GAH1 | 0.60 | 18.82 | 0.69 | 6.81 | 0.67 | 10.40 | 0.89 | 2.05 | 0.89 | 1.98 |
| 2016_IAH2 | 0.80 | 16.25 | 0.29 | 7.05 | 0.38 | 12.27 | 0.92 | 1.63 | 0.93 | 1.50 |
| 2016_IAH3 | 0.26 | 17.48 | 0.31 | 6.25 | 0.16 | 12.23 | 0.63 | 2.76 | 0.60 | 2.34 |
| 2016_IAH4 | 0.58 | 12.91 | 0.75 | 6.09 | 0.76 | 10.67 | 0.90 | 2.12 | 0.92 | 1.58 |
| 2016_INH1 | 0.71 | 18.32 | 0.71 | 5.41 | 0.68 | 8.74 | 0.65 | 3.02 | 0.74 | 2.65 |
| 2016_MIH1 | 0.68 | 17.71 | 0.71 | 6.37 | 0.74 | 9.53 | 0.94 | 1.79 | 0.90 | 2.28 |
| 2016_MOH1 | 0.63 | 16.90 | 0.82 | 3.90 | 0.70 | 8.14 | 0.88 | 2.06 | 0.89 | 1.75 |
| 2016_NEH1 | 0.70 | 13.78 | 0.70 | 3.33 | 0.61 | 11.91 | 0.90 | 0.87 | 0.68 | 1.11 |
| 2016_NEH4 | 0.38 | 15.88 | 0.87 | 2.58 | 0.86 | 7.20 | 0.67 | 1.86 | 0.63 | 1.49 |
| 2016_NYH1 | 0.56 | 13.88 | 0.66 | 8.46 | 0.71 | 11.54 | 0.86 | 2.37 | 0.92 | 1.97 |
| 2016_ONH1 | 0.57 | 12.49 | 0.47 | 10.35 | 0.56 | 11.52 | 0.87 | 2.22 | 0.96 | 1.18 |
| 2016_ONH2 | 0.63 | 14.68 | 0.80 | 4.89 | 0.73 | 10.38 | 0.97 | 1.48 | 0.98 | 1.37 |
| 2016_TXH1 | 0.75 | 12.33 | 0.72 | 4.79 | 0.46 | 17.72 | 0.89 | 1.42 | 0.89 | 1.48 |
| 2016_TXH2 | 0.61 | 16.91 | 0.53 | 6.22 | 0.68 | 8.49 | 0.90 | 1.64 | 0.91 | 1.63 |
| 2016_WIH1 | 0.72 | 14.69 | 0.31 | 6.82 | 0.39 | 11.45 | 0.93 | 1.54 | 0.91 | 1.65 |
| 2016_WIH2 | 0.73 | 19.67 | 0.87 | 3.23 | 0.84 | 5.68 | 0.94 | 1.68 | 0.94 | 1.51 |
| 2017_COH1 | 0.69 | 19.53 | 0.81 | 5.90 | 0.79 | 11.51 | 0.96 | 2.71 | 0.97 | 2.41 |
| 2017_DEH1 | 0.80 | 11.37 | 0.83 | 5.73 | 0.74 | 9.39 | 0.70 | 3.95 | 0.70 | 3.75 |
| 2017_IAH4 | 0.75 | 11.01 | 0.90 | 4.54 | 0.87 | 8.21 | 0.96 | 1.68 | 0.97 | 1.23 |
| 2017_INH1 | 0.68 | 10.55 | 0.88 | 3.24 | 0.77 | 7.58 | 0.85 | 1.96 | 0.87 | 1.80 |
| 2017_MIH1 | 0.58 | 17.12 | 0.69 | 4.32 | 0.68 | 7.70 | 0.71 | 3.72 | 0.76 | 3.75 |
| 2017_MNH1 | 0.31 | 21.25 | 0.32 | 9.63 | 0.33 | 17.57 | 0.81 | 3.25 | 0.82 | 3.22 |
| 2017_MOH1 | 0.42 | 18.09 | 0.92 | 3.67 | 0.87 | 7.55 | 0.91 | 1.64 | 0.92 | 1.46 |
| 2017_NYH1 | 0.63 | 17.58 | 0.74 | 4.68 | 0.79 | 8.83 | 0.93 | 2.07 | 0.92 | 1.84 |
| 2017_NYH2 | 0.39 | 15.32 | 0.76 | 4.70 | 0.80 | 8.01 | 0.94 | 1.86 | 0.95 | 1.66 |
| 2017_NYH3 | 0.48 | 17.60 | 0.94 | 3.63 | 0.94 | 5.42 | 0.86 | 2.21 | 0.87 | 1.89 |
| 2017_ONH1 | 0.55 | 14.23 | 0.58 | 4.80 | 0.12 | 14.09 | 0.82 | 2.35 | 0.83 | 2.22 |
| 2017_ONH2 | 0.54 | 10.98 | 0.45 | 7.45 | 0.57 | 13.30 | 0.74 | 7.69 | 0.71 | 7.17 |
| 2017_TXH1-Dry | 0.80 | 7.93 | 0.78 | 3.10 | 0.68 | 5.59 | 0.92 | 1.32 | 0.92 | 1.30 |
| 2017_TXH1-Early | 0.84 | 6.77 | 0.78 | 3.33 | 0.72 | 6.01 | 0.91 | 1.19 | 0.89 | 1.36 |
| 2017_TXH1-Late | 0.70 | 12.14 | 0.88 | 3.45 | 0.68 | 9.58 | 0.88 | 1.67 | 0.86 | 1.80 |
| 2017_TXH2 | 0.63 | 17.41 | 0.49 | 8.18 | 0.68 | 8.85 | 0.70 | 2.56 | 0.66 | 2.70 |
| 2017_WIH1 | 0.73 | 12.72 | 0.79 | 4.95 | 0.76 | 9.44 | 0.86 | 2.73 | 0.87 | 2.60 |
| 2017_WIH2 | 0.64 | 11.20 | 0.94 | 3.66 | 0.90 | 7.31 | 0.96 | 1.22 | 0.96 | 1.31 |

**Table S6**: Classification into maturity groups and geographical distribution of the 37 environments retained for analysis.

| Environment | Year | Location | Country | Maturity Group |
|---|---|---|---|---|
| 2016_ARH1 | 2016 | ARH1 | USA | Late |
| 2016_ARH2 | 2016 | ARH2 | USA | Late |
| 2016_DEH1 | 2016 | DEH1 | USA | Intermediate |
| 2016_GAH1 | 2016 | GAH1 | USA | Late |
| 2016_IAH2 | 2016 | IAH2 | USA | Early/Intermediate |
| 2016_IAH3 | 2016 | IAH3 | USA | Intermediate/Late |
| 2016_IAH4 | 2016 | IAH4 | USA | Early/Intermediate |
| 2016_INH1 | 2016 | INH1 | USA | Intermediate |
| 2016_MIH1 | 2016 | MIH1 | USA | Early |
| 2016_MOH1 | 2016 | MOH1 | USA | Intermediate |
| 2016_NEH1 | 2016 | NEH1 | USA | Intermediate |
| 2016_NEH4 | 2016 | NEH4 | USA | Intermediate |
| 2016_NYH1 | 2016 | NYH1 | USA | Early |
| 2016_ONH1 | 2016 | ONH1 | Canada | Early |
| 2016_ONH2 | 2016 | ONH2 | Canada | Early |
| 2016_TXH1 | 2016 | TXH1 | USA | Late |
| 2016_TXH2 | 2016 | TXH2 | USA | Late |
| 2016_WIH1 | 2016 | WIH1 | USA | Early/Intermediate |
| 2016_WIH2 | 2016 | WIH2 | USA | Early/Intermediate |
| 2017_COH1 | 2017 | COH1 | USA | Early/Intermediate |
| 2017_DEH1 | 2017 | DEH1 | USA | Intermediate |
| 2017_IAH4 | 2017 | IAH4 | USA | Early/Intermediate |
| 2017_INH1 | 2017 | INH1 | USA | Intermediate |
| 2017_MIH1 | 2017 | MIH1 | USA | Early |
| 2017_MNH1 | 2017 | MNH1 | USA | Early |
| 2017_MOH1 | 2017 | MOH1 | USA | Intermediate |
| 2017_NYH1 | 2017 | NYH1 | USA | Early |
| 2017_NYH2 | 2017 | NYH2 | USA | Early |
| 2017_NYH3 | 2017 | NYH3 | USA | Intermediate |
| 2017_ONH1 | 2017 | ONH1 | Canada | Early |
| 2017_ONH2 | 2017 | ONH2 | Canada | Early |
| 2017_TXH1-Dry | 2017 | TXH1-Dry | USA | Late |
| 2017_TXH1-Early | 2017 | TXH1-Early | USA | Late |
| 2017_TXH1-Late | 2017 | TXH1-Late | USA | Late |
| 2017_TXH2 | 2017 | TXH2 | USA | Late |
| 2017_WIH1 | 2017 | WIH1 | USA | Early/Intermediate |
| 2017_WIH2 | 2017 | WIH2 | USA | Early/Intermediate |

**Table S7**: Estimates of variance components associated with general combining ability (GCA) and specific combining ability (SCA), along wiht standard errors for grain yield (GY, (t $ha^{-1})^2$), plant height (PH, $cm^2$), ear height (EH, $cm^2$), silking (SI, $days^2$), and anthesis (AN, $days^2$), considering all maturity groups combined. Likelihood ratio test (LRT) statistics and *p*-values are reported for each variance component.

| Trait | Class[1] | Effect | Variance Component | Standard Error | LRT-statistic | *p*-value[2] |
|---|---|---|---|---|---|---|
| GY | D | SS | 0.41 | 0.24 | 10.05 | 7.61e-04 |
| | | NSS | 0.92 | 0.31 | 52.65 | 1.99e-13 |
| | | SS × NSS | 0.31 | 0.05 | 192.69 | 2.20e-16 |
| | S | SS | 0.46 | 0.25 | 15.76 | 3.59e-05 |
| | | NSS | 0.88 | 0.32 | 36.83 | 6.43e-10 |
| | | SS × NSS | 0.07 | 0.01 | 194.08 | 2.20e-16 |
| PH | D | SS | 40.07 | 19.63 | 44.58 | 1.22e-11 |
| | | NSS | 135.22 | 40.45 | 120.28 | 2.20e-16 |
| | | SS × NSS | 14.57 | 2.59 | 189.07 | 2.20e-16 |
| | S | SS | 41.92 | 20.07 | 56.21 | 3.26e-14 |
| | | NSS | 130.84 | 39.87 | 94.40 | 2.20e-16 |
| | | SS × NSS | 3.28 | 0.58 | 189.21 | 2.20e-16 |
| EH | D | SS | 21.81 | 10.20 | 59.90 | 4.99e-15 |
| | | NSS | 59.87 | 18.10 | 89.25 | 2.20e-16 |
| | | SS × NSS | 5.83 | 1.31 | 63.88 | 6.66e-16 |
| | S | SS | 21.89 | 10.09 | 73.43 | 2.20e-16 |
| | | NSS | 58.31 | 18.01 | 68.69 | 2.20e-16 |
| | | SS × NSS | 1.30 | 0.29 | 65.67 | 2.78e-16 |
| SI | D | SS | 7.12 | 3.05 | 124.61 | 2.20e-16 |
| | | NSS | 10.80 | 3.07 | 158.46 | 2.20e-16 |
| | | SS × NSS | 0.47 | 0.08 | 192.99 | 2.20e-16 |
| | S | SS | 7.14 | 3.04 | 155.60 | 2.20e-16 |
| | | NSS | 10.85 | 3.11 | 135.30 | 2.20e-16 |
| | | SS × NSS | 0.11 | 0.02 | 192.52 | 2.20e-16 |
| AN | D | SS | 5.73 | 2.48 | 117.25 | 2.20e-16 |
| | | NSS | 9.98 | 2.85 | 154.68 | 2.20e-16 |
| | | SS × NSS | 0.57 | 0.09 | 271.10 | 2.20e-16 |
| | S | SS | 5.67 | 2.43 | 145.55 | 2.20e-16 |
| | | NSS | 10.04 | 2.90 | 126.91 | 2.20e-16 |
| | | SS × NSS | 0.13 | 0.02 | 267.75 | 2.20e-16 |

SS: GCA of Stiff Stalk (SS) lines used as seed parents, reflecting additive effects; NSS: GCA of Non-Stiff Stalk (NSS) lines used as pollen parents, reflecting additive effects; and SS × NSS: SCA of single-cross hybrids between SS and NSS lines, reflecting dominance effects.

[1]This column indicates whether variance components were estimated from models specified with the **D** matrix (Class D) or with the **S** matrix (Class S).

[2]Presented *p*-values are given in scientific notation. All variance components were statistically significant (*p*-value $< 0.001$).

**Table S8**: Estimates of variance components associated with general combining ability (GCA) and specific combining ability (SCA), along wiht standard errors for the early maturity group, focusing on grain yield (GY, (t ha$^{-1}$)$^2$), plant height (PH, cm$^2$), ear height (EH, cm$^2$), silking (SI, days$^2$), and anthesis (AN, days$^2$). Likelihood ratio test (LRT) statistics and $p$-values are reported for each variance component.

| Trait | Class[1] | Effect | Variance Component | Standard Error | LRT-statistic | $p$-value[2] |
|---|---|---|---|---|---|---|
| GY | D | SS | 0.82 | 0.60 | 15.93 | 3.29e-05 |
| | | NSS | 1.49 | 0.76 | 20.51 | 2.97e-06 |
| | | SS × NSS | 0.18 | 0.07 | 46.87 | 3.79e-12 |
| | S | SS | 0.76 | 0.55 | 16.54 | 2.38e-05 |
| | | NSS | 1.42 | 0.73 | 16.93 | 1.94e-05 |
| | | SS × NSS | 0.06 | 0.02 | 51.29 | 4.00e-13 |
| PH | D | SS | 76.58 | 50.86 | 38.33 | 2.98e-10 |
| | | NSS | 106.28 | 49.28 | 44.37 | 1.36e-11 |
| | | SS × NSS | 2.20 | 1.36 | 6.13 | 6.66e-03 |
| | S | SS | 79.46 | 52.60 | 49.02 | 1.27e-12 |
| | | NSS | 107.63 | 49.90 | 42.31 | 3.89e-11 |
| | | SS × NSS | 0.77 | 0.47 | 7.62 | 2.88e-03 |
| EH | D | SS | 22.32 | 15.30 | 25.99 | 1.71e-07 |
| | | NSS | 49.76 | 23.52 | 37.46 | 4.68e-10 |
| | | SS × NSS | 0.71 | 0.81 | 0.93 | 0.17$^{ns}$ |
| | S | SS | 23.05 | 15.64 | 38.60 | 2.61e-10 |
| | | NSS | 50.74 | 23.79 | 36.77 | 6.65e-10 |
| | | SS × NSS | 0.13 | 0.27 | 0.17 | 0.34$^{ns}$ |
| SI | D | SS | 2.60 | 1.99 | 10.69 | 5.38e-04 |
| | | NSS | 6.35 | 3.26 | 19.38 | 5.34e-06 |
| | | SS × NSS | 1.11 | 0.35 | 95.39 | 2.20e-16 |
| | S | SS | 2.93 | 2.15 | 14.57 | 6.75e-05 |
| | | NSS | 6.25 | 3.17 | 17.52 | 1.42e-05 |
| | | SS × NSS | 0.34 | 0.11 | 99.32 | 2.20e-16 |
| AN | D | SS | 2.49 | 1.95 | 9.74 | 9.00e-04 |
| | | NSS | 5.56 | 2.93 | 17.62 | 1.35e-05 |
| | | SS × NSS | 1.29 | 0.39 | 186.69 | 2.20e-16 |
| | S | SS | 2.83 | 2.09 | 15.37 | 4.43e-05 |
| | | NSS | 5.37 | 2.81 | 15.81 | 3.51e-05 |
| | | SS × NSS | 0.40 | 0.12 | 190.60 | 2.20e-16 |

SS: GCA of Stiff Stalk (SS) lines used as seed parents, reflecting additive effects; NSS: GCA of Non-Stiff Stalk (NSS) lines used as pollen parents, reflecting additive effects; and SS × NSS: SCA of single-cross hybrids between SS and NSS lines, reflecting dominance effects.

[1]This column indicates whether variance components were estimated from models specified with the **D** matrix (Class D) or with the **S** matrix (Class S).

[2]Presented $p$-values are given in scientific notation. Variance components showed different levels of significance, including $p < 0.01$ and $p < 0.001$. Cells marked with $^{ns}$ indicate non-significant effects ($p \geq 0.05$).

**Table S9**: Estimates of variance components associated with general combining ability (GCA) and specific combining ability (SCA), along wiht standard errors for the intermediate maturity group, focusing on grain yield (GY, (t $ha^{-1})^2$), plant height (PH, $cm^2$), ear height (EH, $cm^2$), silking (SI, $days^2$), and anthesis (AN, $days^2$). Likelihood ratio test (LRT) statistics and $p$-values are reported for each variance component.

| Trait | Class[1] | Effect | Variance Component | Standard Error | LRT-statistic | $p$-value[2] |
|---|---|---|---|---|---|---|
| GY | D | SS | 0.02 | 0.04 | 0.22 | 0.32$^{ns}$ |
| | | NSS | 0.71 | 0.33 | 24.55 | 3.62e-07 |
| | | SS × NSS | 0.21 | 0.05 | 160.68 | 2.20e-16 |
| | S | SS | 0.03 | 0.05 | 0.66 | 0.21$^{ns}$ |
| | | NSS | 0.30 | 0.23 | 4.77 | 0.02 |
| | | SS × NSS | 0.15 | 0.04 | 177.01 | 2.20e-16 |
| PH | D | SS | 7.19 | 7.12 | 4.22 | 2.00e-02 |
| | | NSS | 145.74 | 61.72 | 53.64 | 1.20e-13 |
| | | SS × NSS | 14.86 | 3.52 | 137.17 | 2.20e-16 |
| | S | SS | 6.73 | 7.13 | 4.92 | 1.32e-02 |
| | | NSS | 113.37 | 55.06 | 13.54 | 1.17e-04 |
| | | SS × NSS | 8.06 | 2.03 | 167.68 | 2.20e-16 |
| EH | D | SS | 23.16 | 17.48 | 30.75 | 1.47e-08 |
| | | NSS | 59.34 | 25.64 | 44.47 | 1.29e-11 |
| | | SS × NSS | 6.83 | 1.80 | 56.91 | 2.28e-14 |
| | S | SS | 21.09 | 16.92 | 11.03 | 4.49e-04 |
| | | NSS | 28.56 | 18.44 | 0.00 | 0.50$^{ns}$ |
| | | SS × NSS | 4.93 | 1.39 | 79.22 | 2.20e-16 |
| SI | D | SS | 3.92 | 2.80 | 138.45 | 2.20e-16 |
| | | NSS | 2.58 | 1.05 | 100.35 | 2.20e-16 |
| | | SS × NSS | 0.07 | 0.02 | 45.59 | 7.29e-12 |
| | S | SS | 3.89 | 2.80 | 90.13 | 2.20e-16 |
| | | NSS | 2.22 | 0.96 | 45.22 | 8.80e-12 |
| | | SS × NSS | 0.06 | 0.02 | 58.35 | 1.09e-14 |
| AN | D | SS | 2.97 | 2.13 | 118.89 | 2.20e-16 |
| | | NSS | 2.41 | 0.98 | 93.55 | 2.20e-16 |
| | | SS × NSS | 0.09 | 0.02 | 74.54 | 2.20e-16 |
| | S | SS | 2.85 | 2.06 | 79.46 | 2.20e-16 |
| | | NSS | 2.05 | 0.89 | 46.96 | 3.62e-12 |
| | | SS × NSS | 0.07 | 0.02 | 89.20 | 2.20e-16 |

SS: GCA of Stiff Stalk (SS) lines used as seed parents, reflecting additive effects; NSS: GCA of Non-Stiff Stalk (NSS) lines used as pollen parents, reflecting additive effects; and SS × NSS: SCA of single-cross hybrids between SS and NSS lines, reflecting dominance effects.

[1]This column indicates whether variance components were estimated from models specified with the **D** matrix (Class D) or with the **S** matrix (Class S).

[2]Presented $p$-values are given in scientific notation. Variance components showed different levels of significance, including $p < 0.05$ and $p < 0.001$. Cells marked with $^{ns}$ indicate non-significant effects ($p \geq 0.05$).

**Table S10**: Estimates of variance components associated with general combining ability (GCA) and specific combining ability (SCA), along wiht standard errors for the late maturity group, focusing on grain yield (GY, $(\text{t ha}^{-1})^2$), plant height (PH, $\text{cm}^2$), ear height (EH, $\text{cm}^2$), silking (SI, $\text{days}^2$), and anthesis (AN, $\text{days}^2$). Likelihood ratio test (LRT) statistics and $p$-values are reported for each variance component.

| Trait | Class[1] | Effect | Variance Component | Standard Error | LRT-statistic | $p$-value[2] |
|---|---|---|---|---|---|---|
| GY | D | SS | 0.11 | 0.09 | 6.40 | 5.71e-03 |
| | | NSS | 0.32 | 0.14 | 59.76 | 5.39e-15 |
| | | SS × NSS | 0.01 | 0.00 | 5.19 | 1.14e-02 |
| | S | SS | 0.11 | 0.09 | 6.38 | 0.01 |
| | | NSS | 0.31 | 0.14 | 40.34 | 1.07e-10 |
| | | SS × NSS | 0.01 | 0.00 | 7.70 | 2.77e-03 |
| PH | D | SS | 7.90 | 6.72 | 7.65 | 2.83e-03 |
| | | NSS | 50.62 | 20.83 | 118.24 | 2.20e-16 |
| | | SS × NSS | 0.42 | 0.22 | 7.60 | 2.92e-03 |
| | S | SS | 7.89 | 6.81 | 7.18 | 3.68e-03 |
| | | NSS | 50.89 | 21.21 | 70.18 | 2.20e-16 |
| | | SS × NSS | 0.40 | 0.22 | 5.68 | 8.58e-03 |
| EH | D | SS | 3.39 | 2.53 | 34.68 | 1.94e-09 |
| | | NSS | 14.37 | 6.18 | 59.34 | 6.66e-15 |
| | | SS × NSS | 0.29 | 0.17 | 4.05 | 2.21e-02 |
| | S | SS | 2.94 | 2.31 | 20.31 | 3.29e-06 |
| | | NSS | 13.88 | 6.33 | 24.93 | 2.97e-07 |
| | | SS × NSS | 0.39 | 0.17 | 13.55 | 1.16e-04 |
| SI | D | SS | 0.75 | 0.55 | 59.23 | 7.05e-15 |
| | | NSS | 2.95 | 1.15 | 113.60 | 2.20e-16 |
| | | SS × NSS | 0.06 | 0.02 | 25.32 | 2.43e-07 |
| | S | SS | 0.80 | 0.59 | 49.05 | 1.25e-12 |
| | | NSS | 2.69 | 1.10 | 69.85 | 2.20e-16 |
| | | SS × NSS | 0.05 | 0.02 | 36.23 | 8.76e-10 |
| AN | D | SS | 0.75 | 0.54 | 66.01 | 2.22e-16 |
| | | NSS | 3.28 | 1.27 | 118.86 | 2.20e-16 |
| | | SS × NSS | 0.08 | 0.02 | 37.19 | 5.35e-10 |
| | S | SS | 0.80 | 0.59 | 48.91 | 1.34e-12 |
| | | NSS | 3.05 | 1.23 | 70.53 | 2.20e-16 |
| | | SS × NSS | 0.06 | 0.02 | 43.59 | 2.03e-11 |

SS: GCA of Stiff Stalk (SS) lines used as seed parents, reflecting additive effects; NSS: GCA of Non-Stiff Stalk (NSS) lines used as pollen parents, reflecting additive effects; and SS × NSS: SCA of single-cross hybrids between SS and NSS lines, reflecting dominance effects.

[1]This column indicates whether variance components were estimated from models specified with the **D** matrix (Class D) or with the **S** matrix (Class S).

[2]Presented $p$-values are given in scientific notation. Variance components showed different levels of significance, including $p < 0.05$, $p < 0.01$, and $p < 0.001$.

**Table S11**: Predictive ability and standard error for grain yield (GY, tons per hectare, t $ha^{-1}$) across training set configurations and prediction methods.

| Training Set Configuration | Method | Predictive Ability | Standard Error |
|---|---|---|---|
| T2 | 1a | 0.735 | 0.031 |
| T2 | 1b | 0.760 | 0.024 |
| T2 | 2a | 0.756 | 0.029 |
| T2 | 2b | 0.780 | 0.021 |
| T2 | 3a | 0.757 | 0.030 |
| T2 | 3b | 0.776 | 0.022 |
| T2 | 4a | 0.768 | 0.022 |
| T2 | 4b | 0.787 | 0.019 |
| T1F | 1a | 0.583 | 0.021 |
| T1F | 1b | 0.562 | 0.018 |
| T1F | 2a | 0.641 | 0.020 |
| T1F | 2b | 0.618 | 0.018 |
| T1F | 3a | 0.608 | 0.020 |
| T1F | 3b | 0.637 | 0.017 |
| T1F | 4a | 0.614 | 0.020 |
| T1F | 4b | 0.636 | 0.017 |
| T1M | 1a | 0.206 | 0.030 |
| T1M | 1b | 0.248 | 0.030 |
| T1M | 2a | 0.244 | 0.030 |
| T1M | 2b | 0.245 | 0.031 |
| T1M | 3a | 0.371 | 0.051 |
| T1M | 3b | 0.411 | 0.036 |
| T1M | 4a | 0.421 | 0.043 |
| T1M | 4b | 0.447 | 0.034 |
| T0 | 1a | -0.241 | 0.020 |
| T0 | 1b | -0.273 | 0.018 |
| T0 | 2a | -0.194 | 0.021 |
| T0 | 2b | -0.272 | 0.018 |
| T0 | 3a | 0.510 | 0.015 |
| T0 | 3b | 0.514 | 0.013 |
| T0 | 4a | 0.512 | 0.013 |
| T0 | 4b | 0.512 | 0.013 |

T2 (both parents of the validation hybrid represented via other crosses), T1F (all hybrids sharing the same seed parent excluded), T1M (all hybrids sharing the same pollen parent excluded), and T0 (no direct parental information in the training set). Methods 1–2 correspond to GBLUP-based multi-kernel models, where method 1 includes only general combining ability (GCA) and method 2 consists of both GCA and specific combining ability (SCA). Methods 3–4 are based on the covariance between tested and untested single-cross hybrids, considering either the additive relationship matrix only (method 3) or both additive and non-additive relationship matrices (method 4). Results are shown for models fitted using either the **D** matrix (a) or the **S** matrix (b).

**Table S12**: Predictive ability and standard error for plant height (PH, centimeters) across training set configurations and prediction methods.

| Training Set Configuration | Method | Predictive Ability | Standard Error |
|---|---|---|---|
| T2 | 1a | 0.855 | 0.009 |
| T2 | 1b | 0.852 | 0.014 |
| T2 | 2a | 0.868 | 0.008 |
| T2 | 2b | 0.864 | 0.013 |
| T2 | 3a | 0.738 | 0.137 |
| T2 | 3b | 0.733 | 0.142 |
| T2 | 4a | 0.734 | 0.113 |
| T2 | 4b | 0.727 | 0.119 |
| T1F | 1a | 0.767 | 0.010 |
| T1F | 1b | 0.766 | 0.009 |
| T1F | 2a | 0.789 | 0.010 |
| T1F | 2b | 0.789 | 0.009 |
| T1F | 3a | 0.415 | 0.019 |
| T1F | 3b | 0.416 | 0.014 |
| T1F | 4a | 0.401 | 0.018 |
| T1F | 4b | 0.406 | 0.014 |
| T1M | 1a | 0.159 | 0.025 |
| T1M | 1b | 0.159 | 0.019 |
| T1M | 2a | 0.170 | 0.025 |
| T1M | 2b | 0.162 | 0.021 |
| T1M | 3a | 0.241 | 0.013 |
| T1M | 3b | 0.239 | 0.018 |
| T1M | 4a | 0.260 | 0.012 |
| T1M | 4b | 0.258 | 0.017 |
| T0 | 1a | -0.100 | 0.022 |
| T0 | 1b | -0.108 | 0.015 |
| T0 | 2a | -0.093 | 0.022 |
| T0 | 2b | -0.106 | 0.015 |
| T0 | 3a | 0.289 | 0.009 |
| T0 | 3b | 0.293 | 0.007 |
| T0 | 4a | 0.288 | 0.008 |
| T0 | 4b | 0.291 | 0.007 |

T2 (both parents of the validation hybrid represented via other crosses), T1F (all hybrids sharing the same seed parent excluded), T1M (all hybrids sharing the same pollen parent excluded), and T0 (no direct parental information in the training set). Methods 1–2 correspond to GBLUP-based multi-kernel models, where method 1 includes only general combining ability (GCA) and method 2 consists of both GCA and specific combining ability (SCA). Methods 3–4 are based on the covariance between tested and untested single-cross hybrids, considering either the additive relationship matrix only (method 3) or both additive and non-additive relationship matrices (method 4). Results are shown for models fitted using either the **D** matrix (a) or the **S** matrix (b).

**Table S13**: Predictive ability and standard error for ear height (EH, centimeters) across training set configurations and prediction methods.

| Training Set Configuration | Method | Predictive Ability | Standard Error |
|---|---|---|---|
| T2 | 1a | 0.805 | 0.013 |
| T2 | 1b | 0.801 | 0.014 |
| T2 | 2a | 0.816 | 0.012 |
| T2 | 2b | 0.816 | 0.014 |
| T2 | 3a | 0.767 | 0.065 |
| T2 | 3b | 0.783 | 0.058 |
| T2 | 4a | 0.758 | 0.056 |
| T2 | 4b | 0.772 | 0.054 |
| T1F | 1a | 0.702 | 0.013 |
| T1F | 1b | 0.700 | 0.013 |
| T1F | 2a | 0.713 | 0.013 |
| T1F | 2b | 0.718 | 0.013 |
| T1F | 3a | 0.296 | 0.012 |
| T1F | 3b | 0.294 | 0.013 |
| T1F | 4a | 0.287 | 0.011 |
| T1F | 4b | 0.287 | 0.013 |
| T1M | 1a | 0.189 | 0.021 |
| T1M | 1b | 0.188 | 0.018 |
| T1M | 2a | 0.204 | 0.020 |
| T1M | 2b | 0.204 | 0.017 |
| T1M | 3a | 0.257 | 0.013 |
| T1M | 3b | 0.265 | 0.015 |
| T1M | 4a | 0.267 | 0.013 |
| T1M | 4b | 0.274 | 0.015 |
| T0 | 1a | -0.141 | 0.023 |
| T0 | 1b | -0.147 | 0.028 |
| T0 | 2a | -0.130 | 0.023 |
| T0 | 2b | -0.128 | 0.028 |
| T0 | 3a | 0.195 | 0.006 |
| T0 | 3b | 0.198 | 0.006 |
| T0 | 4a | 0.192 | 0.005 |
| T0 | 4b | 0.196 | 0.006 |

T2 (both parents of the validation hybrid represented via other crosses), T1F (all hybrids sharing the same seed parent excluded), T1M (all hybrids sharing the same pollen parent excluded), and T0 (no direct parental information in the training set). Methods 1–2 correspond to GBLUP-based multi-kernel models, where method 1 includes only general combining ability (GCA) and method 2 consists of both GCA and specific combining ability (SCA). Methods 3–4 are based on the covariance between tested and untested single-cross hybrids, considering either the additive relationship matrix only (method 3) or both additive and non-additive relationship matrices (method 4). Results are shown for models fitted using either the **D** matrix (a) or the **S** matrix (b).

**Table S14**: Predictive ability and standard error for silking (SI, days) across training set configurations and prediction methods.

| Training Set Configuration | Method | Predictive Ability | Standard Error |
|---|---|---|---|
| T2 | 1a | 0.959 | 0.005 |
| T2 | 1b | 0.961 | 0.006 |
| T2 | 2a | 0.960 | 0.005 |
| T2 | 2b | 0.962 | 0.006 |
| T2 | 3a | 0.912 | 0.075 |
| T2 | 3b | 0.928 | 0.069 |
| T2 | 4a | 0.911 | 0.072 |
| T2 | 4b | 0.926 | 0.066 |
| T1F | 1a | 0.707 | 0.009 |
| T1F | 1b | 0.707 | 0.011 |
| T1F | 2a | 0.721 | 0.009 |
| T1F | 2b | 0.719 | 0.011 |
| T1F | 3a | 0.572 | 0.003 |
| T1F | 3b | 0.572 | 0.004 |
| T1F | 4a | 0.567 | 0.003 |
| T1F | 4b | 0.568 | 0.004 |
| T1M | 1a | 0.646 | 0.021 |
| T1M | 1b | 0.650 | 0.016 |
| T1M | 2a | 0.648 | 0.020 |
| T1M | 2b | 0.648 | 0.015 |
| T1M | 3a | 0.504 | 0.015 |
| T1M | 3b | 0.507 | 0.015 |
| T1M | 4a | 0.517 | 0.015 |
| T1M | 4b | 0.517 | 0.015 |
| T0 | 1a | -0.554 | 0.008 |
| T0 | 1b | -0.554 | 0.008 |
| T0 | 2a | -0.544 | 0.009 |
| T0 | 2b | -0.550 | 0.009 |
| T0 | 3a | 0.538 | 0.004 |
| T0 | 3b | 0.536 | 0.004 |
| T0 | 4a | 0.538 | 0.004 |
| T0 | 4b | 0.536 | 0.004 |

T2 (both parents of the validation hybrid represented via other crosses), T1F (all hybrids sharing the same seed parent excluded), T1M (all hybrids sharing the same pollen parent excluded), and T0 (no direct parental information in the training set). Methods 1–2 correspond to GBLUP-based multi-kernel models, where method 1 includes only general combining ability (GCA) and method 2 consists of both GCA and specific combining ability (SCA). Methods 3–4 are based on the covariance between tested and untested single-cross hybrids, considering either the additive relationship matrix only (method 3) or both additive and non-additive relationship matrices (method 4). Results are shown for models fitted using either the **D** matrix (a) or the **S** matrix (b).

**Table S15**: Predictive ability and standard error for anthesis (AN, days) across training set configurations and prediction methods.

| Training Set Configuration | Method | Predictive Ability | Standard Error |
|---|---|---|---|
| T2 | 1a | 0.950 | 0.008 |
| T2 | 1b | 0.949 | 0.007 |
| T2 | 2a | 0.951 | 0.008 |
| T2 | 2b | 0.950 | 0.007 |
| T2 | 3a | 0.896 | 0.083 |
| T2 | 3b | 0.894 | 0.085 |
| T2 | 4a | 0.894 | 0.077 |
| T2 | 4b | 0.893 | 0.080 |
| T1F | 1a | 0.726 | 0.007 |
| T1F | 1b | 0.722 | 0.011 |
| T1F | 2a | 0.735 | 0.007 |
| T1F | 2b | 0.732 | 0.011 |
| T1F | 3a | 0.536 | 0.004 |
| T1F | 3b | 0.536 | 0.005 |
| T1F | 4a | 0.528 | 0.004 |
| T1F | 4b | 0.529 | 0.005 |
| T1M | 1a | 0.569 | 0.024 |
| T1M | 1b | 0.566 | 0.024 |
| T1M | 2a | 0.571 | 0.023 |
| T1M | 2b | 0.564 | 0.024 |
| T1M | 3a | 0.450 | 0.020 |
| T1M | 3b | 0.436 | 0.020 |
| T1M | 4a | 0.465 | 0.019 |
| T1M | 4b | 0.450 | 0.019 |
| T0 | 1a | -0.482 | 0.011 |
| T0 | 1b | -0.488 | 0.008 |
| T0 | 2a | -0.473 | 0.012 |
| T0 | 2b | -0.486 | 0.008 |
| T0 | 3a | 0.493 | 0.005 |
| T0 | 3b | 0.491 | 0.004 |
| T0 | 4a | 0.493 | 0.005 |
| T0 | 4b | 0.490 | 0.004 |

T2 (both parents of the validation hybrid represented via other crosses), T1F (all hybrids sharing the same seed parent excluded), T1M (all hybrids sharing the same pollen parent excluded), and T0 (no direct parental information in the training set). Methods 1–2 correspond to GBLUP-based multi-kernel models, where method 1 includes only general combining ability (GCA) and method 2 consists of both GCA and specific combining ability (SCA). Methods 3–4 are based on the covariance between tested and untested single-cross hybrids, considering either the additive relationship matrix only (method 3) or both additive and non-additive relationship matrices (method 4). Results are shown for models fitted using either the **D** matrix (a) or the **S** matrix (b).

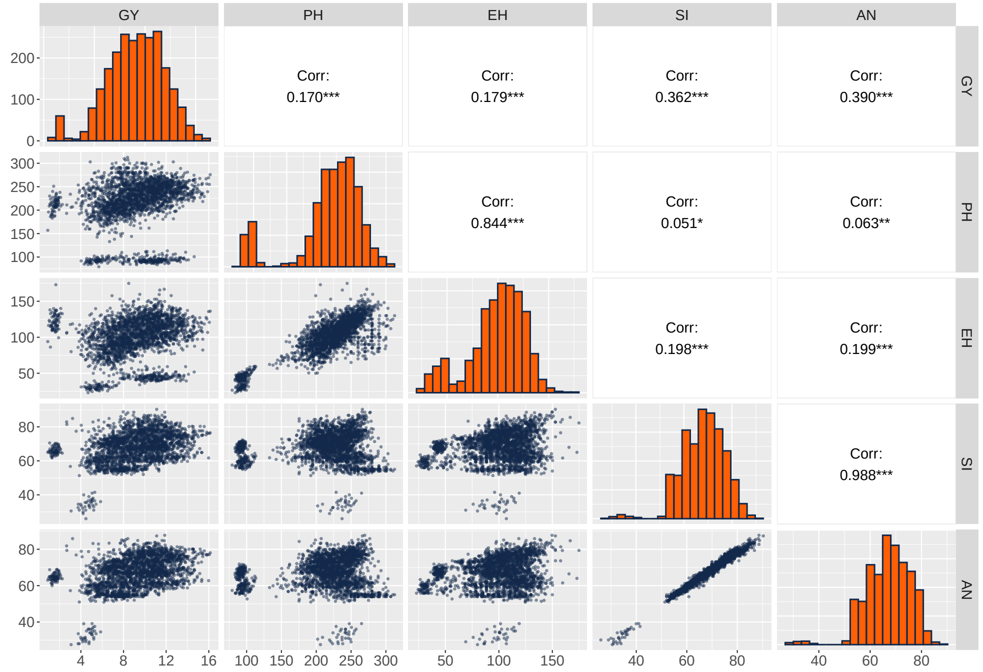


**Fig. S1**: Histograms (diagonal), scatter plots (lower diagonal), phenotypic correlations (upper diagonal), **p*-value < 0.05, ***p*-value < 0.01, and ****p*-value < 0.001. Traits correspond to grain yield (GY, tons per hectare, t ha$^{-1}$), plant height (PH, centimeters), ear height (EH, centimeters), silking (SI, days), and anthesis (AN, days).

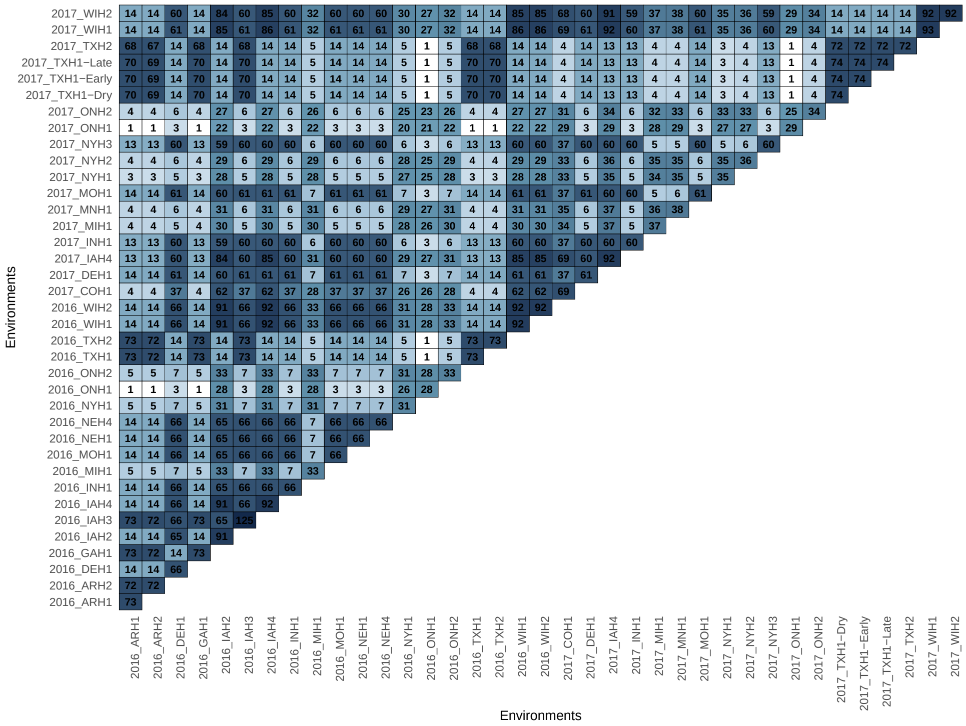

**Fig. S2**: Co-occurrence matrix of hybrids among environments. The diagonal values represent the number of unique hybrids in a given environment, whereas off-diagonal values represent the number of unique common hybrids among environments.

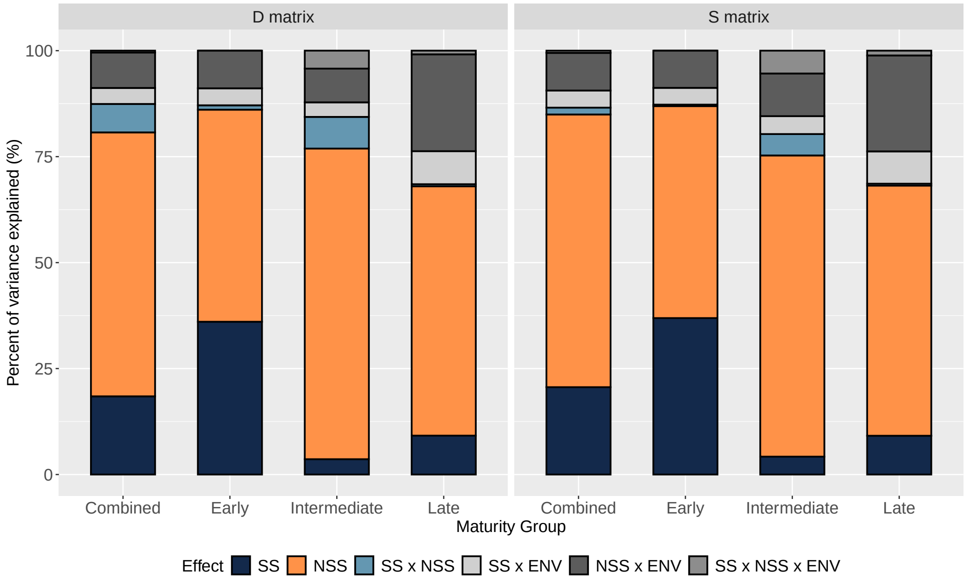


**Fig. S3**: Percent of variance explained for plant height (PH, centimeters). Variance components were estimated from models specified with either the **D** matrix or the **S** matrix. SS: general combining ability (GCA) of Stiff Stalk (SS) lines used as seed parents, reflecting additive effects; NSS: GCA of Non-Stiff Stalk (NSS) lines used as pollen parents, reflecting additive effects; SS × NSS: specific combining ability (SCA) of single-cross hybrids between SS and NSS lines, reflecting dominance effects; SS × ENV: interaction between SS lines and environments; NSS × ENV: interaction between NSS lines and environments; SS × NSS × ENV: three-way interaction among SS, NSS, and environments. Results are shown for all hybrids (Combined) as well as separately by maturity group (Early, Intermediate, Late).

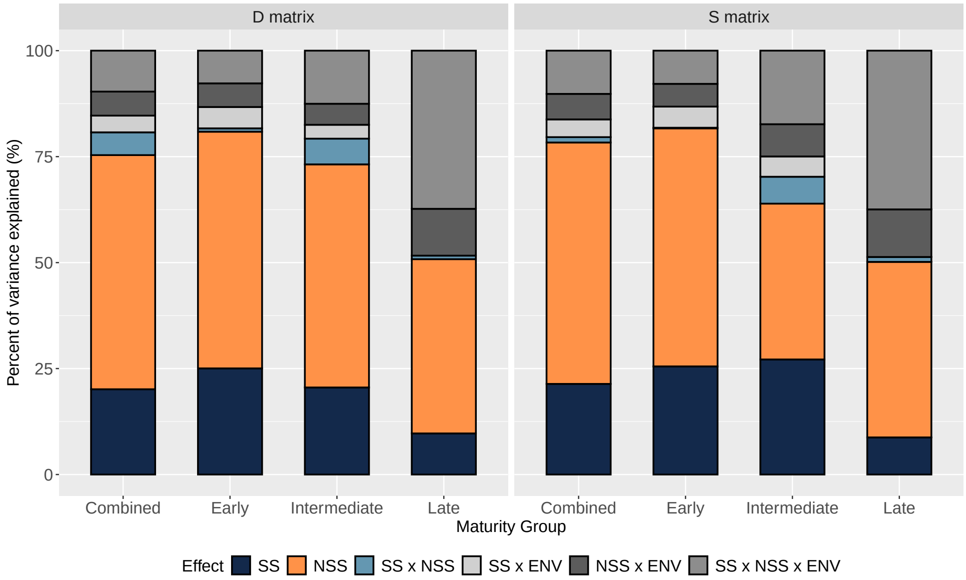


**Fig. S4**: Percent of variance explained for ear height (EH, centimeters). Variance components were estimated from models specified with either the **D** matrix or the **S** matrix. SS: general combining ability (GCA) of Stiff Stalk (SS) lines used as seed parents, reflecting additive effects; NSS: GCA of Non-Stiff Stalk (NSS) lines used as pollen parents, reflecting additive effects; SS × NSS: specific combining ability (SCA) of single-cross hybrids between SS and NSS lines, reflecting dominance effects; SS × ENV: interaction between SS lines and environments; NSS × ENV: interaction between NSS lines and environments; SS × NSS × ENV: three-way interaction among SS, NSS, and environments. Results are shown for all hybrids (Combined) as well as separately by maturity group (Early, Intermediate, Late).

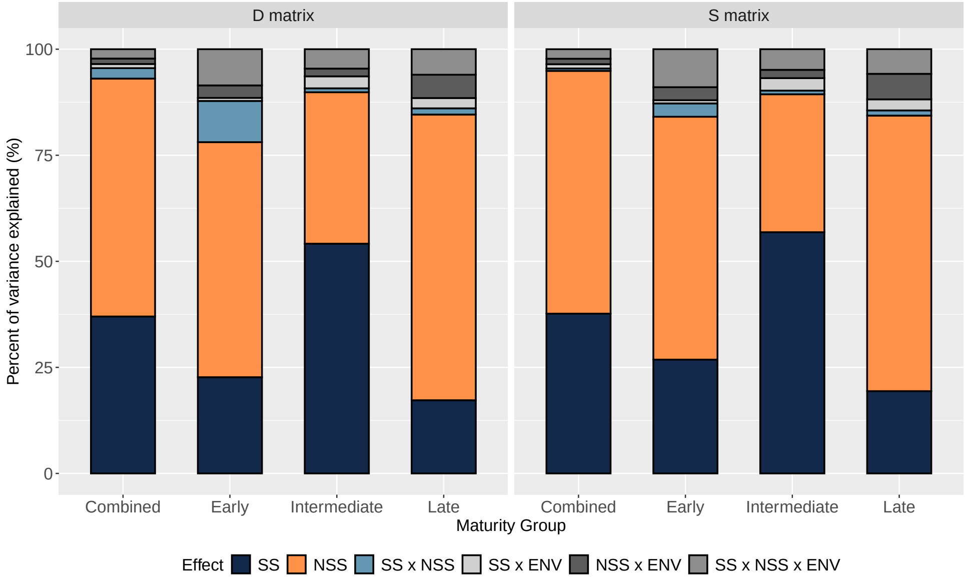


**Fig. S5**: Percent of variance explained for silking (SI, days). Variance components were estimated from models specified with either the **D** matrix or the **S** matrix. SS: general combining ability (GCA) of Stiff Stalk (SS) lines used as seed parents, reflecting additive effects; NSS: GCA of Non-Stiff Stalk (NSS) lines used as pollen parents, reflecting additive effects; SS × NSS: specific combining ability (SCA) of single-cross hybrids between SS and NSS lines, reflecting dominance effects; SS × ENV: interaction between SS lines and environments; NSS × ENV: interaction between NSS lines and environments; SS × NSS × ENV: three-way interaction among SS, NSS, and environments. Results are shown for all hybrids (Combined) as well as separately by maturity group (Early, Intermediate, Late).

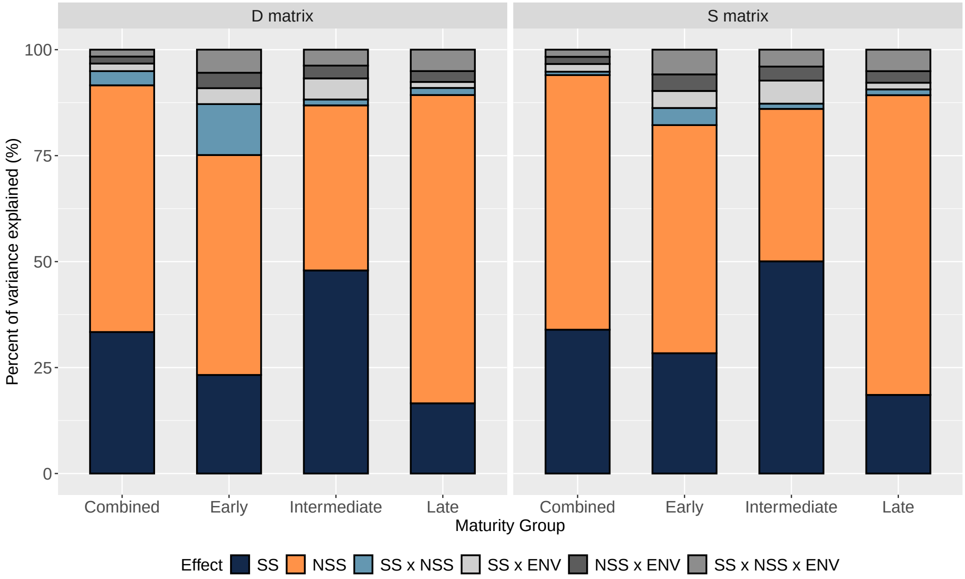


**Fig. S6**: Percent of variance explained for anthesis (AN, days). Variance components were estimated from models specified with either the **D** matrix or the **S** matrix. SS: general combining ability (GCA) of Stiff Stalk (SS) lines used as seed parents, reflecting additive effects; NSS: GCA of Non-Stiff Stalk (NSS) lines used as pollen parents, reflecting additive effects; SS $\times$ NSS: specific combining ability (SCA) of single-cross hybrids between SS and NSS lines, reflecting dominance effects; SS $\times$ ENV: interaction between SS lines and environments; NSS $\times$ ENV: interaction between NSS lines and environments; SS $\times$ NSS $\times$ ENV: three-way interaction among SS, NSS, and environments. Results are shown for all hybrids (Combined) as well as separately by maturity group (Early, Intermediate, Late).
